# *Yersinia* Type III-Secreted Effectors Evade the Caspase-4 Inflammasome in Human Cells

**DOI:** 10.1101/2023.01.24.525473

**Authors:** Jenna Zhang, Igor E. Brodsky, Sunny Shin

## Abstract

*Yersinia* are gram-negative zoonotic bacteria that use a type III secretion system (T3SS) to inject *Yersinia* outer proteins (Yops) into the host cytosol to subvert essential components of innate immune signaling. However, *Yersinia* virulence activities can elicit activation of inflammasomes, which lead to inflammatory cell death and cytokine release to contain infection. *Yersinia* activation and evasion of inflammasomes have been characterized in murine macrophages but remain poorly defined in human cells, particularly intestinal epithelial cells (IECs), a primary site of intestinal *Yersinia* infection. In contrast to murine macrophages, we find that in both human IECs and macrophages, *Yersinia pseudotuberculosis* T3SS effectors enable evasion of the caspase-4 inflammasome, which senses cytosolic lipopolysaccharide (LPS). The antiphagocytic YopE and YopH, as well as the translocation regulator YopK, were collectively responsible for evading inflammasome activation, in part by inhibiting *Yersinia* internalization mediated by YadA and β1-integrin signaling. These data provide insight into the mechanisms of *Yersinia-*mediated inflammasome activation and evasion in human cells, and reveal species-specific differences underlying regulation of inflammasome responses to *Yersinia*.

**Importance:** *Yersinia* are responsible for significant disease burdens in humans, ranging from recurrent disease outbreaks (yersiniosis) to pandemics (*Yersinia pestis* plague). Together with rising antibiotic resistance rates, there is a critical need to better understand *Yersinia* pathogenesis and host immune mechanisms, as this information will aid in developing improved immunomodulatory therapeutics. Inflammasome responses in human cells are less studied relative to murine models of infection, though recent studies have uncovered key differences in inflammasome responses between mice and humans. Here, we dissect human intestinal epithelial cell and macrophage inflammasome responses to *Yersinia pseudotuberculosis.* Our findings provide insight into species- and cell type-specific differences in inflammasome responses to *Yersinia*.

## Introduction

Innate immunity is a critical component of host defense and employs pattern recognition receptors (PRRs) to detect pathogen-associated molecular patterns (PAMPs) (1, 2). Specific cytosolic PRRs in the Nucleotide-binding domain and Leucine-rich Repeat-containing protein (NLR) family induce the formation and activation of multimeric immune complexes known as inflammasomes in response to cytosolic PAMPs and virulence-associated activities (3–5). The NLRP3 inflammasome is activated by a variety of stimuli, including potassium efflux downstream of bacterial-induced pore formation (5–9), whereas the NAIP-NLRC4 inflammasome responds to bacterial flagellin and components of bacterial type III secretion systems (T3SS) (10–18). Inflammasomes recruit and activate caspase-1, which in turn processes members of the IL-1 family of cytokines and the pore-forming protein GSDMD into their active forms (19–23). In addition to these canonical inflammasome pathways, a non-canonical caspase-4 inflammasome in humans, and orthologous caspase-11 inflammasome in mice, responds to cytosolic lipopolysaccharide (LPS) during gram-negative bacterial infection to initiate GSDMD processing, pore formation, and cytokine release (24–31). Consequently, inflammasome activation leads to release of active IL-1 family cytokines, specifically IL-1β and IL-18, and an inflammatory form of cell death known as pyroptosis, that collectively amplify immune signaling and promote anti-bacterial defense (32). In parallel, bacterial infection triggers other programmed cell death pathways, including apoptosis and necroptosis, thereby providing multiple layers of defense against bacterial pathogens (33, 34).

The enteric pathogenic Yersiniae, *Yersinia pseudotuberculosis* and *Y. enterocolitica,* express a conserved T3SS that injects *Yersinia* outer proteins (Yops) into target cells (35). T3SS-injected Yops manipulate host cellular pathways to promote infection, but also activate and inhibit effector-triggered cell death responses. For example, in murine macrophages, YopJ-mediated suppression of NF-κB signaling induces apoptosis, whereas YopE-mediated disruption of cytoskeletal dynamics activates the pyrin inflammasome and hyper-translocation of T3SS components in the absence of YopK activates the NLRP3 and caspase-11 inflammasomes (36–45). Notably, *Yersinia* utilizes two other effectors, YopM and YopK, to evade YopE-triggered pyrin and T3SS-triggered NLRP3/caspase-11-inflammasome activation respectively and promote infection (42, 43, 45, 46). Collectively, these virulence strategies manipulate the host in order to facilitate colonization and evade ensuing effector-triggered immunity (47).

Inflammasome responses to *Yersinia* have primarily been studied in murine macrophages. However, fundamental differences exist between human and murine inflammasome responses to bacterial infection that may impact pathogenesis and disease severity in the host, including ligand recognition, virulence-driven immune suppression, and inflammasome component expression (48–68). Furthermore, beyond phagocytic cells, inflammasomes are expressed in multiple cell types, including intestinal epithelial cells (IECs) (69), which are a primary site of infection and interact significantly with enteric pathogens during intestinal colonization. While specialized IECs known as M cells, which overlie intestinal immune compartments called Peyer’s patches, are considered the primary point of bacterial invasion of the intestinal epithelial barrier (70)(71)(72)(73), *Yersinia* microcolonies are frequently found outside of Peyer’s patches within submucosal pyogranulomas and in close proximity to non-M cell IECs (74). Importantly, many enteric bacterial pathogens use secreted virulence factors to inhibit IEC death in order to preserve their replicative niche during early infection (75, 76). *Y. enterocolitica* uses two anti-phagocytic Yops, YopE and YopH, to evade caspase-1 and NLRP3-dependent inflammasome activation in human IECs (77). However, the role of other Yops or other inflammasomes in IECs and human cells broadly is unknown.

Here, we find that in contrast to prior findings in murine macrophages (38, 44, 78, 79), the *Y. pseudotuberculosis* (*Yptb*) effector YopJ does not induce death of human IECs or macrophages. In contrast, in human cells, *Yptb* evades caspase-4- and GSDMD-dependent cell death and inflammasome-dependent cytokine release. Notably, the type III secreted effectors YopE, YopH, and YopK, collectively enable *Yptb* to evade caspase-4 activation in human cells. Mechanistically, YopE and YopH blockade of bacterial internalization prevented accumulation of intracellular bacteria and correlated with reduced inflammasome activation. Furthermore, the *Yptb* adhesin YadA and host β1-integrins were required for *Yptb* internalization into human IECs and subsequent caspase-4 activation in the absence of YopE, YopH, and YopK. These findings demonstrate a key role for disruption of actin-mediated internalization in *Yersinia* evasion of the noncanonical inflammasome in human IECs, and uncover important species and cell-type-specific differences in inflammasome responses to *Yersinia* infection.

## Results

### *Yptb* effectors evade T3SS-dependent inflammasome activation in human cells

During infection of murine macrophages, *Yersinia* injects Yops to evade inflammasome activation (42, 43, 45, 46). However, wild type (WT) *Yersinia* induces rapid cell death (Fig. 1A) and caspase-8 and caspase-1 activation due to YopJ-mediated inhibition of NF-κB and MAPK signaling, enabling antibacterial defense despite effector-mediated immune modulation (38, 44, 78, 79). In contrast, human macrophages and IECs do not undergo YopJ-dependent cell death early during *Y. enterocolitica* infection or at all during *Y. pseudotuberculosis* infection (56, 77, 80). In agreement with these findings, infection of the human colorectal cell line Caco-2 (Fig. 1B) or primary human monocyte-derived macrophages (hMDMs) (Fig. 1C) with WT *Yptb* failed to induce release of lactate dehydrogenase (LDH), a marker of cell lysis, into the supernatant. Furthermore, WT *Yptb* infection of the human monocytic cell line THP-1 macrophages similarly failed to induce LDH release (Fig. S1A), collectively suggesting that in contrast to murine cells, human cells do not undergo cell death in response to WT *Yptb* infection.

**Figure 1.**
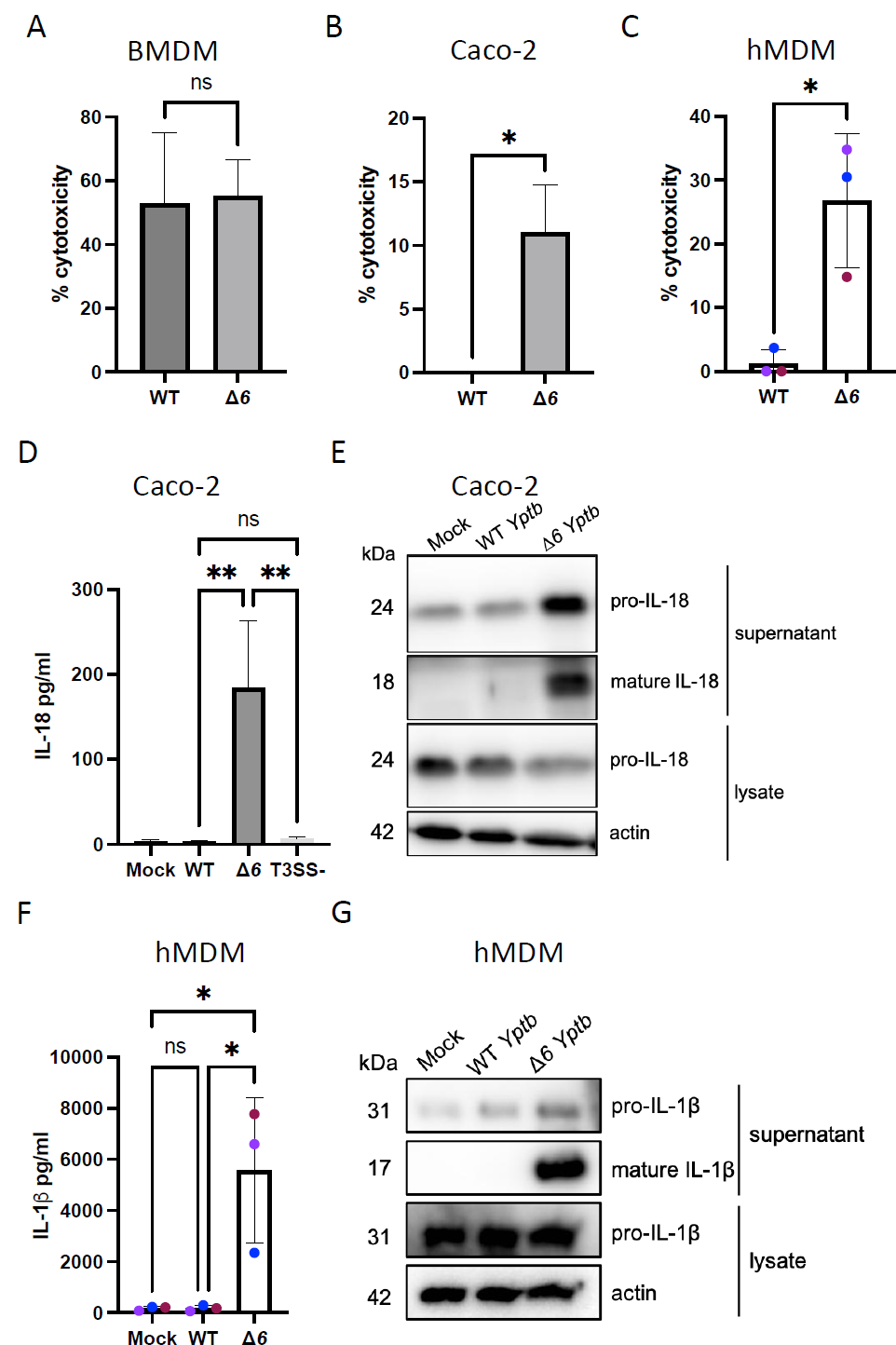
*Yptb* effectors evade T3SS-dependent inflammasome activation in human cells. BMDMs (A), Caco-2 cells (B, D-E) or hMDMs (C, F-G) were infected with PBS (Mock), WT *Yptb*, or Δ*6 Yptb*. (A-C) Cell death was measured as percent cytotoxicity normalized to cells treated with 2% Triton at 6hpi. (D) Release of IL-18 into the supernatant was measured by ELISA at 6hpi. (E) Lysates and supernatants collected 6hpi were immunoblotted for IL-18 and β-actin. (F) Release of IL-1β into the supernatant was measured by ELISA at 6hpi. (G) Lysates and supernatants collected 6hpi were immunoblotted for IL-1β and β-actin. ns — not significant, * p < 0.05, ** p < 0.01 by t-test (A-C) or one-way ANOVA (D, F). Shown are averages and error bars representing the standard deviation from at least three pooled experiments.

Lack of cell death in *Yptb-*infected human cells could be due to a failure to initiate programmed cell death pathways, or due to active evasion or suppression of programmed cell death by the bacteria. *Y. enterocolitica* deploys two injected effectors, YopH and YopE, to evade inflammasome activation in human IECs (77). Critically, *Yptb* lacking all six of its injected effectors (Δ*6 Yptb*) triggered significantly higher levels of cell death compared to either mock or WT *Yptb*-infected cells in Caco-2 IECs (Fig. 1B), hMDMs (Fig. 1C) and THP-1 macrophages (Fig. S1A). Further, Δ*6 Yptb*-infected polarized and non-polarized Caco-2 cells robustly cleaved and released the inflammasome-dependent cytokine IL-18 compared to mock or WT *Yptb*-infected cells (Fig. S1B, Fig. 1D-E). IL-18 release following Δ*6 Yptb* infection occurred at a range of increasing doses, whereas WT *Yptb* failed to induce IL-18 release even at the highest doses (Fig. S1C). Consistent with our and others’ findings that expression and release of the inflammasome-dependent cytokine IL-1β is very low in human IECs (48, 49, 77), we did not detect IL-1β release during Δ*6 Yptb* infection (Fig. S1D). While IECs do not produce IL-1β, this cytokine is released by macrophages undergoing pyroptosis. Indeed, hMDMs and THP-1 macrophages infected with Δ*6 Yptb* robustly cleaved and secreted IL-1β in contrast to mock or WT *Yptb*-infected cells (Fig. 1F-G, Fig. S1E), collectively indicating that in the absence of its injected effectors, *Yersinia* triggers inflammasome activation in multiple human cell types. As expected, inflammasome activation in response to Δ*6 Yptb* infection is T3SS-dependent, as an isogenic *Yptb* strain cured of its virulence plasmid encoding the T3SS did not induce IL-18 release in human IECs (Fig. 1D). Collectively, these results indicate that Yops enable WT *Yptb* to evade inflammasome activation in human cells, a fundamentally distinct outcome from the induction of apoptosis and pyroptosis triggered in murine macrophages by WT *Yersinia* (38, 44, 78, 79).

### Caspase-4 is required for Δ*6 Yptb*-induced inflammasome activation in human cells

Caspases play important roles in cleaving and activating inflammasome-dependent cytokines and executing cell death (34). Pretreatment of Caco-2 cells with the pan-caspase inhibitor ZVAD prior to infection with Δ*6 Yptb* completely abrogated IL-18 release and cell death compared to the vehicle control (Fig. 2A-B), indicating that caspases mediate inflammasome activation downstream of Δ*6 Yptb* infection in human IECs. As ZVAD broadly inhibits multiple caspases, we next sought to determine which specific caspases are required for Δ*6 Yptb*-induced inflammasome activation in human IECs. Caspase-1 is critical for inflammasome-dependent cytokine release and pyroptosis during bacterial infection (34, 44, 64, 81), and is activated during *Y. enterocolitica* infection of Caco-2 cells (77). Δ*6 Yptb* infection of two independent clones of *CASP1^-/-^* Caco-2 cells (49) resulted in a partial loss of IL-18 release compared to WT Caco-2 cells (Fig. S2A). Similarly, WT Caco-2s pretreated with the caspase-1 inhibitor YVAD exhibited incomplete loss of IL-18 release after Δ*6 Yptb* infection compared to DMSO vehicle-treated cells (Fig. S2B), suggesting that caspase-1 contributes to, but is not absolutely required for, Δ*6 Yptb*-induced inflammasome activation. Caspase-8 is activated in murine macrophages and IECs in response to infection by multiple pathogens, including *Yersinia* (44, 82, 83), and can process caspase-1 substrates such as IL-1β and GSDMD to mediate pyroptosis in the absence of caspase-1 (64, 82, 84). Notably, siRNA knockdown of *CASP8* and pretreatment with the caspase-8 inhibitor IETD resulted in a partial reduction in IL-18 release following Δ*6 Yptb* infection (Fig. S2C-E), suggesting that like caspase-1, caspase-8 partially contributes to inflammasome activation during Δ*6 Yptb* infection. However, *CASP8* siRNA knockdown in *CASP1*^-/-^ Caco-2 cells did not abrogate inflammasome activation during Δ*6 Yptb* infection (Fig. S2F-G), indicating that additional caspase activation pathways likely mediate cell death during Δ*6 Yptb* infection of human IECs.

**Figure 2.**
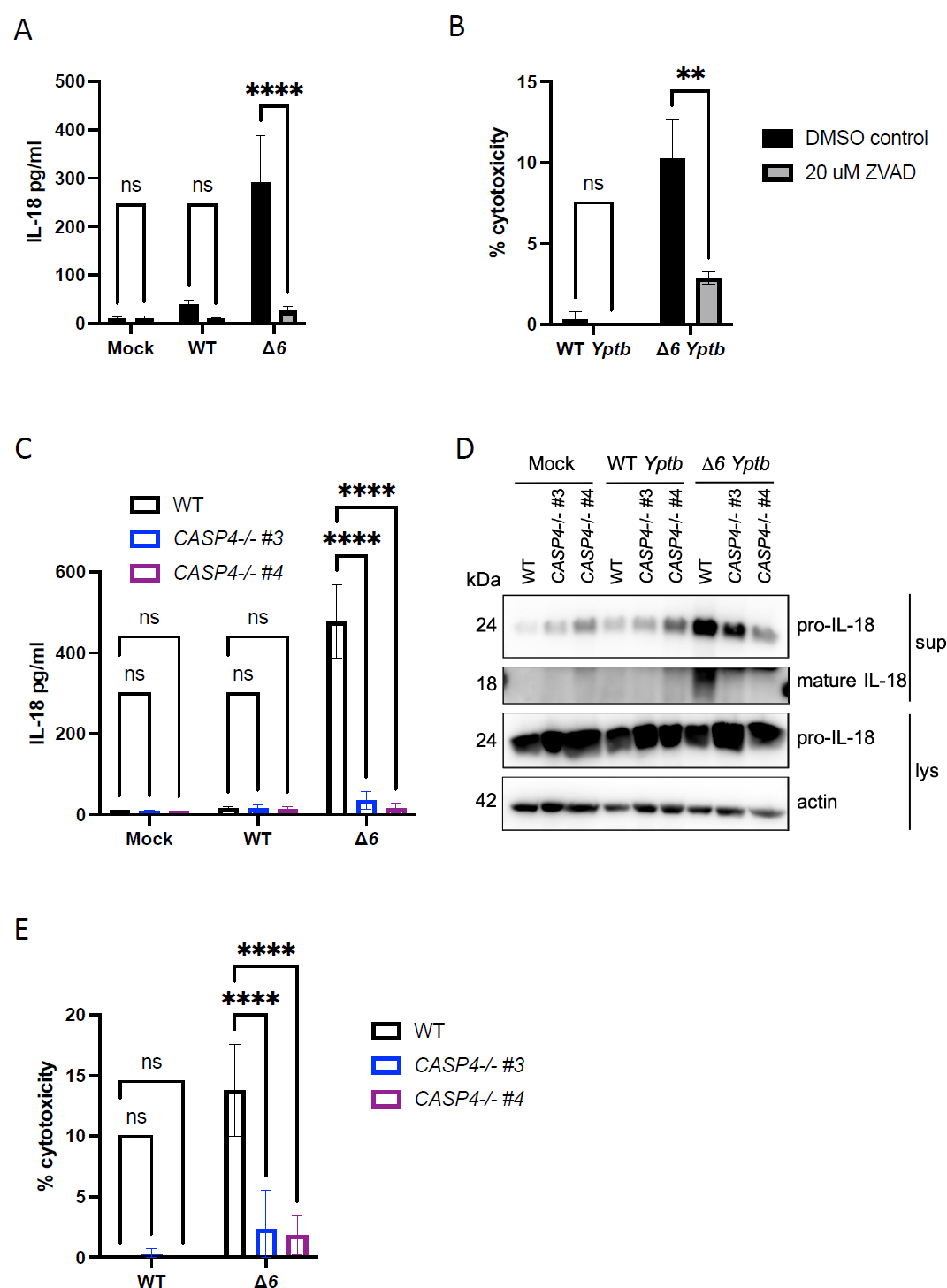
Caspase-4 is required for Δ*6 Yptb*-induced inflammasome activation in human cells. (A, B) One hour prior to infection, WT Caco-2 cells were treated with 20 μM ZVAD or DMSO as a vehicle control. Cells were then infected with PBS (Mock), WT *Yptb* or Δ*6 Yptb*. At 6hpi (A) release of IL-18 into the supernatant was measured by ELISA and (B) cell death was measured as percent cytotoxicity normalized to cells treated with 2% triton. (C-E) WT or two independent single cell clones of *CASP4*^−/−^ Caco-2 cells were infected with PBS (Mock), WT *Yptb* or Δ*6 Yptb*. (C) Release of IL-18 into the supernatant was measured by ELISA at 6hpi. (D) Lysates and supernatants collected 6hpi were immunoblotted for IL-18 and β-actin. (E) Cell death was measured at 6hpi as percent cytotoxicity normalized to cells treated with 2% triton. ** p < 0.01, **** p < 0.0001 by two-way ANOVA. Shown are averages and error bars representing the standard deviation from at least three pooled experiments.

Caspase-4 plays a critical role in inflammasome responses to a variety of enteric pathogens in human IECs (48, 49, 60, 67), and its activation triggers both IEC death and IL-18 release (48). To test whether caspase-4 contributes to inflammasome responses to Δ*6 Yptb*, we infected two independent single-cell clones of *CASP4*^−/−^ Caco-2 cells (49) with either WT or Δ*6 Yptb*. As expected, WT *Yptb* infection did not elicit inflammasome activation in either WT or *CASP4*^−/−^ Caco-2 cells, and Δ*6 Yptb* infection of WT Caco-2 cells resulted in robust release of cleaved IL-18 and cell death. Notably, *CASP4* deficiency abrogated cleavage and release of active IL-18 and cell death in response to Δ*6 Yptb*, indicating that caspase-4 is absolutely required for inflammasome responses to Δ*6 Yptb* infection in human IECs (Fig. 2C-E). Further, *CASP4* deficiency in THP-1 macrophages (85) resulted in a partial but significant decrease in inflammasome activation (Fig. S3A), suggesting that Yop-mediated evasion of the caspase-4 inflammasome is a conserved evasion mechanism across human cell types. Loss of inflammasome activation in THP-1 macrophages lacking caspase-4 largely mirrored our findings with pretreatment with the pan-caspase inhibitor ZVAD (Fig. S3B), further supporting caspase-4 contribution to inflammasome activation in human macrophages. Caspase-5 contributes to *Salmonella*-induced inflammasome activation in Caco-2 cells (49). To test whether caspase-5 also contributes to Δ*6 Yptb*-induced inflammasome activation in Caco-2 cells, we treated WT Caco-2 cells with either a control scramble siRNA or *CASP5* siRNA. Knockdown of *CASP5* resulted in a partial decrease in IL-18 release (Fig. S3C and S3D), suggesting that caspase-5 contributes but is not absolutely required for inflammasome activation. Collectively, these data indicate for the first time that *Yptb* deploys its Yops to evade caspase-4 inflammasome activation in human macrophages and IECs.

### GSDMD is required for Δ*6 Yptb*-induced inflammasome-dependent cytokine release and cell death in human cells

Inflammasome activation leads to cleavage of the protein GSDMD and liberation of its active pore-forming N-terminal domain, leading to its oligomerization into a large ungated pore (19, 25, 86). Formation of the GSDMD pore in the plasma membrane leads to release of IL-1 family cytokines as well as cell lysis and death, collectively termed “pyroptosis” (87–90). Caspase-4 cleaves and activates GSDMD via release of its N-terminal domain (25, 91). Notably, Δ*6 Yptb* infection led to robust GSDMD cleavage in WT Caco-2 cells, which was completely absent in *CASP4*^−/−^ Caco-2 cells, indicating that caspase-4 is required for GSDMD cleavage in human IECs in response to *Yersinia* lacking its secreted effectors (Fig. 3A). In contrast, consistent with a lack of observed cell death and IL-18 release, WT *Yptb* infection did not elicit GSDMD cleavage in either WT or *CASP4*^−/−^ IECs (Fig. 3A). To test whether GSDMD is required for cell death and inflammasome-dependent cytokine release during Δ*6 Yptb* infection, we pretreated Caco-2 cells with disulfiram, a chemical inhibitor of GSDMD pore formation (92). Critically, disulfiram treatment completely abrogated IL-18 release and cell death downstream of inflammasome activation in Δ*6 Yptb*-infected cells compared to infected vehicle control treated cells (Fig. 3B-C). Consistent with this observation in human IECs, disulfiram treatment of human THP-1 macrophages resulted in an abrogation of IL-1β release (Fig. 3D). Collectively, these results indicate that *Yptb* Yops enable evasion of the caspase-4 inflammasome and GSDMD activation.

**Figure 3.**
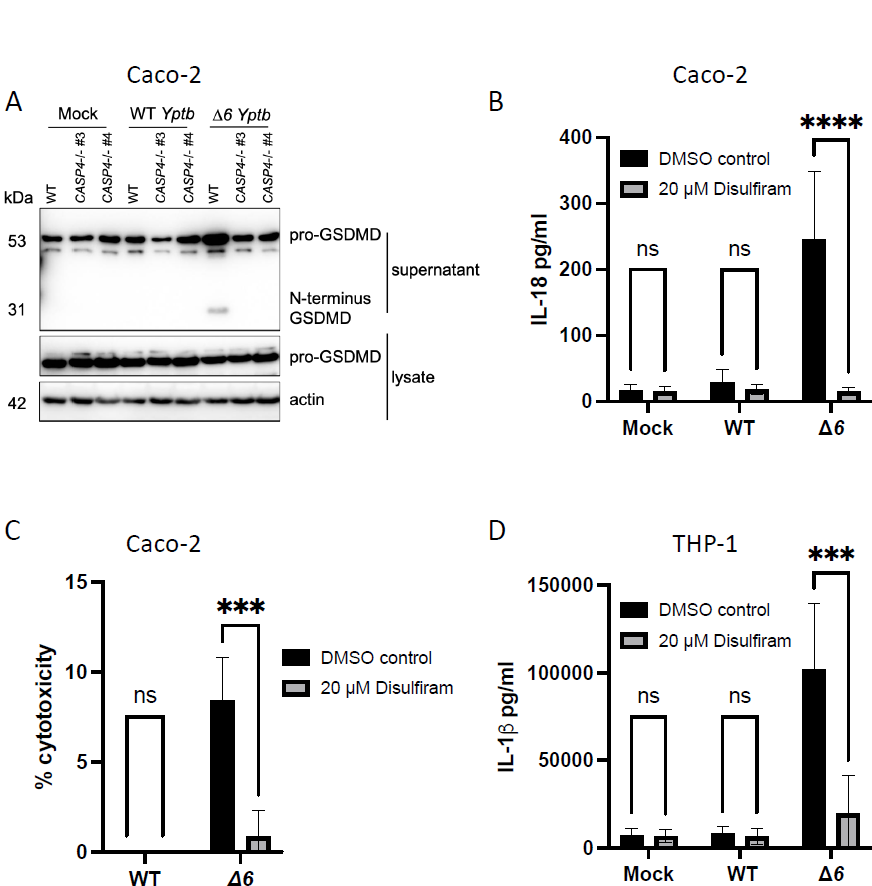
GSDMD is required for Δ*6 Yptb*-induced inflammasome-dependent cytokine release and cell death in human cells. (A) WT or two independent single cell clones of *CASP4*^−/−^ Caco-2 cells were infected with PBS (Mock), WT *Yptb* or Δ*6 Yptb*. Lysates and supernatants were collected at 6hpi and immunoblotted for GSDMD and β-actin. One hour prior to infection, (B, C) WT Caco-2 cells or (D) THP-1 macrophages were treated with 30 μM disulfiram or DMSO as a vehicle control. Cells were then infected with PBS (Mock), WT *Yptb* or Δ*6 Yptb*. Release of (B) IL-18 or (D) IL-1β into the supernatant and (C) percent cytotoxicity normalized to cells treated with 2% triton were measured at 6hpi. ***p<0.001, **** p < 0.0001 by two-way ANOVA. Shown are averages and error bars representing the standard deviation from at least three pooled experiments.

The NLRP3 inflammasome can be activated by a variety of stimuli during infection, including potassium efflux downstream of caspase-4-dependent GSDMD activation and pore formation (5–9). Previous studies of human IEC responses during *Y. enterocolitica* infection identified a critical role for the NLRP3 inflammasome (77). However, studies of human IECs found that NLRP3 does not play a role in inflammasome activation in response to *Salmonella* infection, potentially due to very low levels of NLRP3 expression in human IECs as compared to human macrophages (49, 93, 94). In agreement, WT Caco-2 cells pretreated with a chemical inhibitor of the NLRP3 inflammasome, MCC950, underwent comparable levels of inflammasome activation in response to Δ*6 Yptb* infection as infected vehicle control-treated Caco-2 cells (Fig. S4A). Caco-2 cells stimulated with LPS and nigericin, a known agonist of the NLRP3 inflammasome, also failed to induce IL-18 release, further suggesting a lack of NLRP3 inflammasome activity in Caco-2 cells. The NAIP/NLRC4 inflammasome, which senses and responds to flagellin and T3SS ligands (10–18) (Fig. S4B), and the inflammasome adaptor protein ASC (Fig. S4C) were also dispensable for Δ*6 Yptb*-induced inflammasome activation, consistent with prior findings that expression of these proteins is very low in human IECs (49). Collectively, these results indicate that the canonical NLRP3 and NAIP/NLRC4 inflammasomes, as well as broadly ASC-dependent inflammasomes, are not activated during Δ*6 Yptb* infection of Caco2 cells, and that GSDMD cleavage and activation occurs downstream of caspase-4 and is required for cell death and cytokine release.

### YopE, YopH, and YopK synergistically enable *Yptb* to evade human inflammasome responses

Our findings demonstrate that *Yptb* lacking its entire repertoire of injected effectors induce inflammasome activation in human IECs (Fig. 1). In contrast, Caco-2 cells infected with a panel of *Yptb* mutant strains each lacking one of the six Yops failed to elicit IL-18 secretion (Fig. 4A), indicating that loss of any single secreted Yop was insufficient to alleviate inflammasome evasion and that several Yops likely have overlapping functions in evading inflammasome activation. Notably, single loss of YopK and YopM failed to induce inflammasome activation (Fig. 4A), despite their roles in evading the NLRP3/caspase-11 and pyrin inflammasomes respectively in murine macrophages (40, 43, 45, 46). *Y. enterocolitica* was previously reported to regulate NLRP3 inflammasome activation in Caco-2 cells by a combination of YopE and YopH-mediated blockade of integrin signaling (77). Despite a lack of a role for the NLRP3 inflammasome during Δ*6 Yptb* infection of human IECs (Fig. S4A), *Yptb* lacking both YopE and YopH (Δ*yopEH Yptb*) elicited significantly elevated IL-18 release in Caco-2 cells, indicating that combinatorial loss of both YopE and YopH was sufficient to induce inflammasome activation in human IECs (Fig. 4B). Nonetheless, IL-18 levels during Δ*yopEH Yptb* infection were significantly lower than IL-18 levels released during Δ*6 Yptb* infection (Fig. 4B), suggesting that additional Yops contribute to inflammasome evasion during *Yersinia* infection of human IECs.

**Figure 4.**
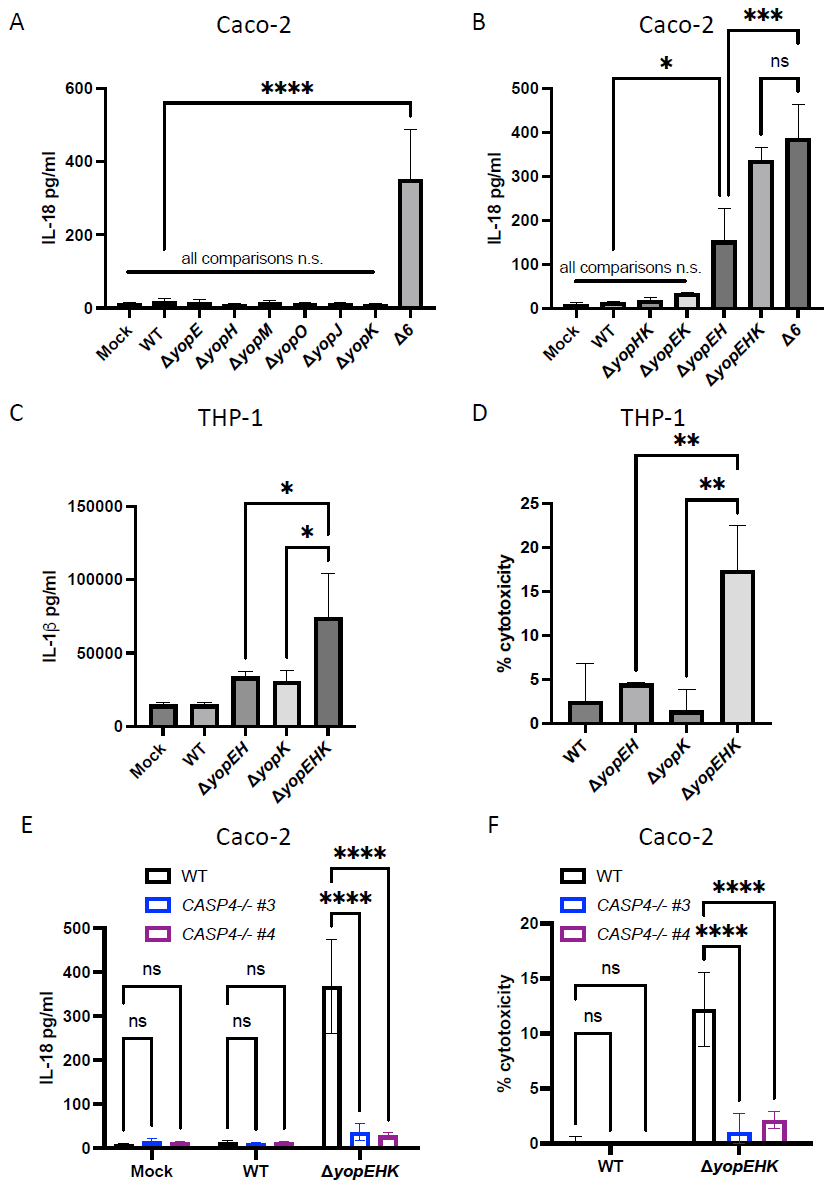
YopE, YopH, and YopK synergistically enable *Yptb* to evade human inflammasome responses. (A, B) WT Caco-2 cells or (C, D) WT THP-1 macrophages were infected with PBS (Mock) or indicated strain of *Yptb*. Release of (A, B) IL-18 or (C) IL-1β into the supernatant and (D) percent cytotoxicity normalized to cells treated with 2% triton was measured at 6hpi. (E, F) WT or two independent single cell clones of *CASP4*^−/−^ Caco-2 cells were infected with PBS (Mock) or the indicated strain of *Yptb*. (E) Release of IL-18 into the supernatant and (F) percent cytotoxicity normalized to cells treated with 2% triton were measured at 6hpi. * p < 0.05, ** p < 0.01, *** p < 0.001, **** p < 0.0001 by one-way ANOVA (A-D) or two-way ANOVA (E, F). Shown are averages and error bars representing the standard deviation from at least three pooled experiments.

YopK is a translocated effector that negatively regulates the translocation of other Yops and T3SS components (42, 43, 45). In murine macrophages, YopK evades inflammasome activation, whereas YopE and YopH do not contribute to inflammasome evasion (42, 43, 45). Given that deletion of *yopK* alone failed to elicit IL-18 release in human IECs (77) (Fig. 4A), we considered that YopK could modulate inflammasome activation in human IECs in a manner that is masked by YopE and YopH, perhaps because these effectors are hypertranslocated in a Δ*yopK* mutant. Indeed, IL-18 release during Δ*yopEHK Yptb* infection was substantially elevated compared to Δ*yopEH Yptb,* and fully recapitulated levels of IL-18 release observed during Δ*6 Yptb* infection (Fig. 4B), indicating that YopE, YopH and YopK function together to enable *Yptb* evasion of inflammasome activation during infection. Critically, individual loss of YopK, dual loss of YopK and YopE (Δ*yopEK*) or dual loss of YopK and YopH (Δ*yopHK*) all failed to induce inflammasome activation. Only in a *yopEH* mutant background did additional deletion of YopK lead to an increase in IL-18 release (Fig. 4B). Consistent with infection of human IECs, Δ*yopEHK* infection also triggered higher levels of inflammasome activation than either Δ*yopEH* or Δ*yopK* infection in human THP-1 macrophages (Fig. 4C-D). Further, as with Δ*6 Yptb* infection, Δ*yopEHK*-induced inflammasome activation in Caco-2 cells was fully dependent on caspase-4 (Fig. 4E-F). Taken together, these data suggest that YopE, YopH, and YopK act in concert to evade the noncanonical inflammasome in both human IECs and macrophages. These results are distinct from murine macrophages, in which YopE instead activates the pyrin inflammasome, and YopK and YopM contribute to inflammasome evasion (42, 43, 45, 46).

### YopE and YopH inhibit caspase-4-dependent inflammasome activation in human IECs by blocking actin-dependent bacterial internalization

During *Y. enterocolitica* infection of human IECs, YopE and YopH evade NLRP3 inflammasome activation by disrupting IL-18 transcriptional priming downstream of invasin-β1-integrin signaling (77). However, *Yersinia* adhesin-β1-integrin-mediated signaling also triggers host cytoskeletal rearrangements to facilitate bacterial internalization (95–98). YopE and YopH disruption of focal adhesion complexes and actin filamentation consequently inhibits *Yersinia* uptake into host cells (98–108). Notably, consistent with both Δ*6* and Δ*yopEHK Yptb* infection, we found that caspase-4 was absolutely required for inflammasome activation induced by Δ*yopEH Yptb* (Fig. 5A and 5B). Considering YopE and YopH’s known roles in inhibiting bacterial internalization, we hypothesized that YopE- and YopH-mediated inflammasome evasion in human IECs might be linked to their antiphagocytic activity, which would limit bacterial internalization, subsequent cytosolic delivery of LPS, and caspase-4 inflammasome activation. Consistent with previous reports (98, 100–102, 104, 108), levels of *Yptb* internalization into human cells 2 hours post-infection were lowest in WT *Yptb*-infected cells, while Δ*yopEH*-infected cells had significantly elevated levels of intracellular bacteria, as assessed by colony forming units (CFUs) (Fig. S5A-B). Δ*yopEH* and Δ*6*-infected cells had comparable levels of intracellular bacteria at 2 hpi, indicating that YopE and YopH regulate internalization into human cells, while YopK limits inflammasome activation through a mechanism distinct from bacterial internalization. In agreement with our CFU data, microscopic analysis of Caco-2 cells infected with GFP-expressing bacteria demonstrated that Δ*yopEH* and Δ*6 Yptb*-infected cells had significantly higher levels of intracellular bacteria (i.e. GFP-only bacteria) than WT *Yptb*-infected cells (Fig. S5C-D). Furthermore, coinfection of Caco-2 cells with Δ*6 Yptb* and increasing doses of WT *Yptb* resulted in reduced bacterial internalization (Fig. 5C), which corresponded to a dose-dependent decrease in inflammasome activation (Fig. 5D) as compared to Δ*6 Yptb* alone infected cells. These results suggest that WT *Yptb*-injected Yops act in trans and can complement inhibition of bacterial uptake and inflammasome activation during Δ*6 Yptb* infection. As YopE and YopH block bacterial uptake via disruption of the actin cytoskeleton (98, 101–107), we asked whether pharmacological inhibition of the actin cytoskeleton would prevent inflammasome activation in response to Δ*yopEH Yptb* infection. Notably, cytochalasin D, an inhibitor of actin polymerization previously demonstrated to block bacterial internalization (102, 109), significantly reduced both intracellular bacterial numbers and inflammasome activation during Δ*yopEH* infection (Fig. S5E-F). Collectively, these results highlight a role for YopE and YopH-mediated internalization inhibition in contributing to inflammasome evasion in human IECs.

**Figure 5.**
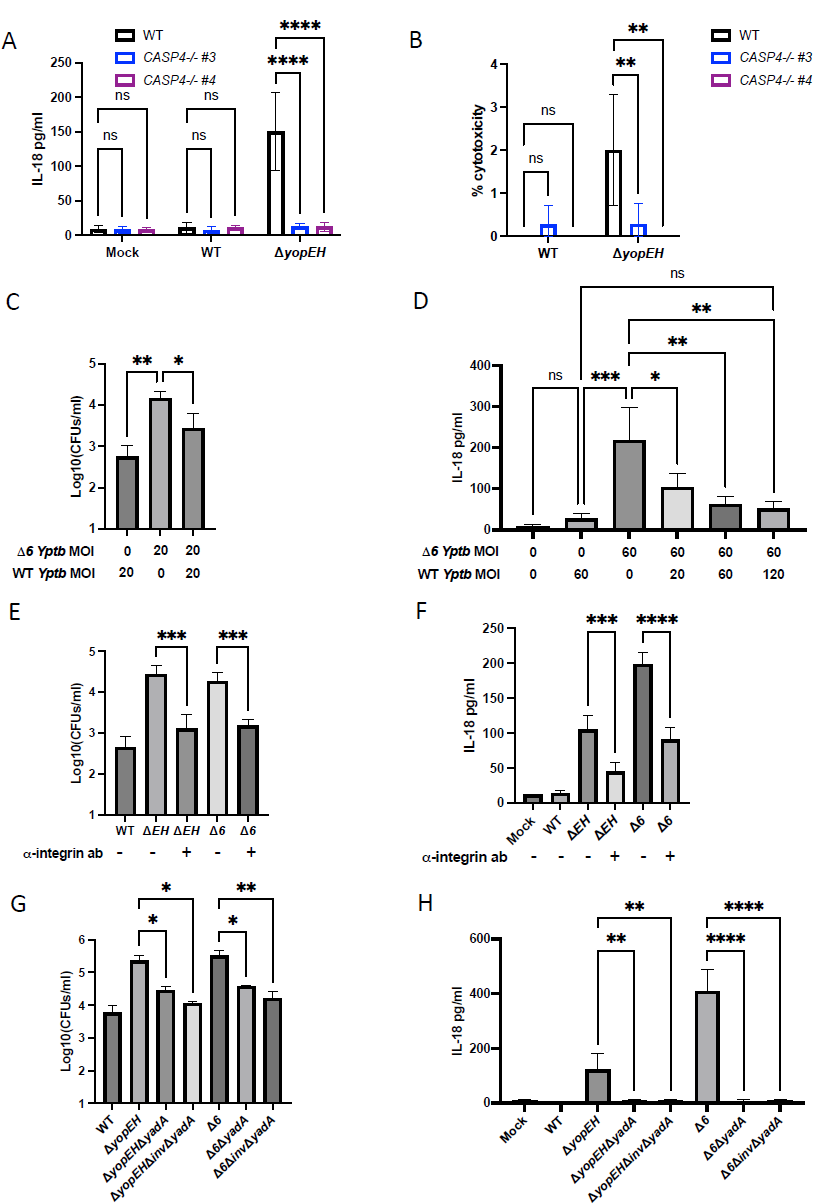
YopE and YopH evade caspase-4-dependent inflammasome activation and actin-dependent bacterial internalization in human IECs. (A, B) WT or two independent single cell clones of *CASP4*^−/−^ Caco-2 cells were infected with PBS (Mock) or the indicated strain of *Yptb*. (A) Release of IL-18 into the supernatant and (B) percent cytotoxicity normalized to cells treated with 2% triton were measured at 6hpi. (C, D) WT Caco-2 cells were infected with the indicated MOI and strain of *Yptb*. (C) Cells were lysed at 2 hpi and bacteria were plated on *Yersinia*-selective agar to calculate CFUs. (D) Release of IL-18 into the supernatant was measured at 6hpi. (E, F) WT Caco-2 cells were incubated for one hour with or without a 1:50 dilution of antibody against β-1 integrin and then infected with PBS (Mock) or the indicated strain of *Yptb* at an MOI of 20 (E) or 60 (F). (E) Cells were lysed at 2 hpi and bacteria were plated on *Yersinia*-selective agar to calculate CFUs. (F) Release of IL-18 into the supernatant was measured at 6hpi. (G, H) WT Caco-2 cells were infected with PBS (Mock) or the strain of *Yptb* at an MOI of 20 (G) or 60 (H). (G) Cells were lysed at 2 hpi and bacteria were plated on *Yersinia*-selective agar to calculate CFUs. (H) Release of IL-18 into the supernatant was measured at 6hpi. * p < 0.05, ** p < 0.01, *** p < 0.001, **** p < 0.0001 by two-way ANOVA (A, B) or one-way ANOVA (C-H). Shown are averages and error bars representing the standard deviation from at least three pooled experiments.

### YadA and β1-integrin are necessary for bacterial internalization and inflammasome activation

Because Cytochalasin D has pleiotropic inhibitory effects, we sought to evaluate the role of bacterial internalization in triggering inflammasome activation by manipulating components of β1-integrin signaling. As macrophages are naturally phagocytic and do not require β1-integrin signaling to induce bacterial uptake, we focused on human IECs. *Yersinia*-β1-integrin interactions initiate an intracellular signaling cascade, including phosphorylation of the serine/threonine kinase Akt, that ultimately regulates actin-dependent cellular changes necessary for bacterial internalization (110–115). Consistent with previous reports, chemical inhibition of Akt isoforms 1 and 2 diminished Δ*6 Yptb* internalization in IECs (Fig. S6A) (116). Critically, inflammasome activation induced by Δ*6 Yptb* was also significantly blunted in the presence of Akt inhibitors (Fig. S6B). Further, pretreatment with an anti-β1-integrin blocking antibody previously found to reduce *Yersinia* entry into cells (117), also significantly decreased bacterial internalization and inflammasome activation during both Δ*yopEH* and Δ*6 Yptb* infection (Fig. 5E-F). Moreover, siRNA knockdown of α5β1 integrin similarly reduced levels of internalized bacteria and inflammasome activation (Fig. S6C-E). Together, these data suggest that disruption of β1-integrin receptor engagement and downstream components of β1-integrin signaling prevents *Yersinia* internalization and limits inflammasome activation in IECs.

The *Yersinia* adhesin, invasin, binds to β1-integrin to facilitate efficient bacterial internalization into mammalian cells, particularly during early intestinal invasion and translocation through M cells (70, 118). Inflammasome activation during *Y. enterocolitica* infection of human IECs was found to be dependent on invasin-β1-integrin signaling that upregulates *Il18* transcript levels (77). However, unlike *Ye, Yptb* does not express high levels of invasin under T3SS-expressing conditions (117, 119, 120). Consistently, deletion of invasin had a minor impact on Δ*yopEH* and Δ*6 Yptb*-induced internalization and inflammasome activation (Fig. S6F-G) suggesting the possibility that other adhesins may mediate T3SS-dependent inflammasome activation by *Yptb*. YadA is encoded on the *Yersinia* virulence plasmid, and its expression is co-regulated with the T3SS and Yops (121). As YadA mediates efficient cellular entry under conditions where invasin expression is repressed (117), we hypothesized that YadA-mediated bacterial internalization might drive caspase-4-dependent inflammasome activation. Indeed, deletion of YadA in both Δ*yopEH* and Δ*6 Yptb* significantly reduced bacterial internalization to levels comparable to WT infection (Fig. 5G) and completely abrogated inflammasome activation (Fig. 5H). Notably, deletion of YadA alone largely phenocopied deletion of both invasin and YadA with respect to internalization and inflammasome activation (Fig. 5G-H), implicating YadA as the primary adhesin responsible for bacterial internalization and inflammasome activation during *Yptb* infection of human IECs.

## Discussion

In this study, we report for the first time that the *Yptb* T3SS-injected effectors YopE, YopH and YopK collectively enable evasion of caspase-4 inflammasome activation and pyroptosis in human cells (Fig. 1-4, S1, S3). Mechanistically, YopE and YopH prevented bacterial internalization and, together with YopK, limited caspase-4 inflammasome activation downstream of YadA-β1-integrin-mediated uptake of *Yptb* into IECs (Fig. 5, S5-6).

The NLRP3 inflammasome is reported to be activated by *Y. enterocolitica* in human IECs (77). However, we found that NLRP3 was dispensable for inflammasome responses to *Yptb* in Caco-2 cells (Fig. S4A). NLRP3 expression is very low in Caco-2 IECs and primary human epithelium (49, 93, 94), and other enteric pathogens and known NLRP3 stimuli fail to activate the NLRP3 inflammasome in Caco-2 cells (49) (Fig. S4A). Differences in cell culture conditions or between *Y. enterocolitica* and *Y. pseudotuberculosis* may account for the differential role of NLRP3 during infection of human IECs. Moreover, the adaptor protein ASC, which is necessary for function of the NLRP3, pyrin, and other inflammasomes, was dispensable for inflammasome responses to Δ*6 Yptb* (Fig. S4C), indicating that canonical inflammasomes are unlikely to be involved in this response. Notably, although YopE-mediated cytoskeletal disruption activates the pyrin inflammasome, which is inhibited by YopM during infection of murine macrophages (40, 46), YopM had no impact on inflammasome activation in human IECs even in the presence of YopE (Fig. 4A), potentially due in part to low expression of pyrin in IECs (122, 123).

We found instead for the first time that *Yptb* activates the caspase-4 inflammasome in human epithelial cells (Fig. 2C-E, 4E-F, 5A-B). Caspase-4 is more highly expressed in human IECs than other inflammasome components (49), and a broad range of intracellular enteric pathogens engage or inhibit caspase-4/11 in human and murine IECs (48, 49, 76, 60, 62–67, 75). We also found a partial role for caspase-1 and caspase-8 in inflammasome activation in Caco-2 cells (Fig. S2A-B, S2D-E). Caspase-8 and caspase-1 can be recruited to the same inflammasome complexes and have compensatory or sequential roles (44, 64, 81).

Simultaneous knockdown of caspase-1 and caspase-8 still allowed for release of IL-18 during Δ*6 Yptb* infection (Fig. S2F), suggesting that they are unlikely to play overlapping roles and may act sequentially. Whether caspase-8 and capase-1 are activated downstream of caspase-4 or are acting in a parallel pathway during *Yersinia* infection of human IECs is unknown. CASP4^−/−^ THP-1 macrophages also had reduced levels of inflammasome activation during Δ*6 Yptb* infection (Fig S3A-B), suggesting that like in human IECs, caspase-4 contributes to Δ*6 Yptb*-induced inflammasome activation in human macrophages.

In addition to blocking entry into epithelial cells (98, 101–104) (Fig. S5), YopE and H also block bacterial internalization into M cells (77), specialized follicular epithelial cells overlying Peyer’s patches that are considered the primary site of entry across the intestinal epithelium for *Yersinia* (70–73, 104). As such, invasin-dependent entry into M cells may occur prior to full T3SS upregulation, thereby allowing the bacteria to cross the epithelium without triggering inflammasome activation. Intriguingly, we found that ι1*6 Yptb* induces inflammasome activation during both apical and basolateral infection of polarized Caco-2 cells, suggesting a role for non-M cell IECs in inflammasome-mediated host defense. While integrins are predominantly expressed apically on M cells and basolaterally on non-M cell IECs, intestinal pathogens can interact with basolateral host receptors at cellular junctions either due to depolarization events leading to apical receptor relocation or to focal discontinuities leading to luminal exposure of basolateral elements (124–126). Furthermore, *Yptb* microcolonies are found outside Peyer’s patches within submucosal pyogranulomas in close proximity basolaterally to IECs (74). YopE and YopH may prevent apical or basolateral uptake by non M-cell IECs that would otherwise trigger inflammasome-mediated responses within the intestinal epithelium.

*Y. entercolitica* YopE and YopH were previously reported to inhibit inflammasome activation by blocking a priming signal downstream of integrin signaling that upregulates *IL18* transcript levels (77). However, integrin signaling is also crucial for bacterial internalization into IECs. Using a variety of orthogonal approaches, we show here that disruption of integrin signaling reduced inflammasome activation commensurate with reduced levels of *Yptb* internalization by IECs (Fig. 5, S6). Additionally, YadA, rather than invasin, was required for *Δ6 Yptb-*induced inflammasome activation and bacterial internalization (Fig. 5G-H), consistent with *Yptb* downregulation of invasin and YadA co-expression with the T3SS (117, 121). Our findings lead us to propose a model in which YopE/H-dependent blockade of YadA-mediated bacterial uptake by IECs limits delivery of LPS into the host cell, thereby allowing bacteria to evade the noncanonical inflammasome. Because adhesin-integrin binding and signal transduction mediate a variety of cellular outcomes, it is difficult to fully exclude the impact of additional factors on inflammasome activation and evasion facilitated by YopE and YopH.

While YopE and YopH interfere with inflammasome responses during *Y. enterocolitica* infection of human IECs, no role was previously found for the *Y. enterocolitica* homolog of YopK, YopQ (77). Consistently, while loss of YopK alone had no effect on inflammasome responses to *Yptb*, combinatorial loss of YopK, YopE, and YopH recapitulated the levels of inflammasome activation observed with Δ*6 Yptb* infection, suggesting that YopK evades a component of an inflammasome pathway masked by YopE and YopH (Fig. 4). In murine macrophages, YopK prevents LPS-mediated inflammasome activation by preventing destabilization of the *Yersinia*-containing vacuole (42, 43). In the absence of YopE, YopH, and YopK, *Yersinia* may be more readily taken up and exposed to the host cell cytosol, potentially due to vacuolar damage as a result of T3SS-mediated pore formation (42, 127). We found that loss of YopK augments Δ*yopEH*-induced inflammasome activation in both human macrophages and IECs, potentially pointing to a conserved mechanism by which YopK, E, and H enable *Yptb* to evade inflammasome responses across human cell types. Further studies are needed to determine precisely how the T3SS and YopK evade cytosolic *Yersinia* LPS exposure and caspase-4 activation.

The synergistic evasion of the caspase-4 inflammasome by YopE, YopH, and YopK, as well as the absence of YopJ-induced cell death, in both human IECs and THP-1 macrophages is fundamentally distinct from the response to *Yptb* observed in murine macrophages. Human cells may be intrinsically more resistant to pathogen-induced cell death due to altered expression of prosurvival factors such as cellular FLICE-inhibitory protein (cFLIP) or A20.

Furthermore, unlike macrophages, IECs are not phagocytic, potentially allowing contamination of the host cell cytosol by bacterial LPS to be a more specific indicator of virulence activity in IECs and providing a rationale for increased reliance on caspase-4-mediated responses. Further studies dissecting inflammasome responses between murine and human IECs and macrophages will provide additional insight into cell type- and species-specific differences in the mechanisms of inflammasome activation by bacterial pathogens.

Overall, our data demonstrate that Yops E, H, and K, enable *Yersinia pseudotuberculosis* to evade caspase-4 inflammasome responses downstream of YadA-β1-integrin signaling in human cells, thereby revealing a major difference in interactions between Yops and inflammasomes in murine and human macrophages. Our study further highlights the distinct nature of inflammasome responses and bacterial effector activities in different cell types in mice and humans, which provides insight into how inflammasome responses and bacterial virulence activities shape health and disease in mice and humans.

## Materials and Methods

### Ethics statement

All studies involving primary human monocyte-derived macrophages (hMDMs) were performed in compliance with the requirements of the US Department of Health and Human Services and the principles expressed in the Declaration of Helsinki. hMDMs from de-identified healthy human donors were obtained from the University of Pennsylvania Human Immunology Core, which holds Institutional Review Board approval. These samples are considered a secondary use of de-identified human specimens and are exempt via Title 55 Part 46, Subpart A of 46.101 (b) of the Code of Federal Regulations.

### Bacterial strains and growth conditions

*Yersinia* strains are described in Table S1 in the supplemental material. Δ*yopEHK*, Δ*yopEK*, and Δ*yopHK* were generated by introducing a frameshift mutation of the *yopK* open reading frame into the Δ*yopEH*, Δ*yopE* and Δ*yopH* backgrounds respectively using a plasmid provided by Dr. James Bliska and an allelic exchange method (128). Δ*yopM* was generated by introducing an unmarked deletion of the *yopM* open reading frame into IP2666 using a plasmid provided by Dr. James Bliska and the same allelic exchange method. Δ*yopEH*Δ*inv* and Δ*6*Δ*inv* strains were generated by introducing an unmarked deletion of the *invasin* open reading frame into Δ*yopEH* and Δ*6* strains respectively using a plasmid provided by Dr. Joan Mecsas and the same allelic exchange method. Δ*yopEH*Δ*yadA*, Δ*yopEH*Δ*inv*Δ*yadA*, Δ*6*Δ*yadA*, and Δ*6*Δ*inv*Δ*yadA* strains were generated by introducing a kanamycin resistance cassette in place of the *yadA* open reading frame into Δ*yopEH,* Δ*yopEH*Δ*inv,* Δ*6,* and Δ*6*Δ*inv* strains respectively using a plasmid provided by Dr. Petra Dersch and the same allelic exchange method. Yersiniae were cultured overnight at 26°C with aeration in 2x yeast extract-tryptone (YT) broth. To induce T3SS expression, in the morning, the bacteria were diluted into fresh 2xYT containing 20 mM sodium oxalate and 20 mM MgCl_2_. Bacteria were grown with aeration for 1 hour at 26°C followed by 2 hour at 37°C prior to infection. All cultures were pelleted at 6000 x *g* for 3 min and resuspended in phosphate-buffered saline (PBS). Cells were infected at an MOI of 60 unless otherwise indicated, centrifuged at 290 x *g* for 10 min and incubated at 37°C. At 1 hour post-infection, epithelial cells were treated with 20 ng/ml or 100 ng/ml of gentamicin for 6 hour or 2 hour time points respectively and macrophages were treated with 100 ng/ml of gentamicin for all time points. Infections proceeded at 37°C for the indicated length of time for each experiment. In all experiments control cells were mock infected with PBS.

### Cell culture conditions

All cells were grown at 37°C in a humidified incubator with 5% CO_2_.

#### Cell culture of Caco-2 cells

Caco-2 cells (HTB-27; American Type Culture Collection) were maintained in Dulbecco’s modified Eagle’s medium (DMEM) supplemented with 10% (vol/vol) heat-inactivated fetal bovine serum (FBS), 100 IU/mL penicillin and 100 μg/mL streptomycin. One day prior to infection, Caco-2 cells were incubated with 0.25% trypsin-EDTA (Gibco) diluted 1:1 with 1 x PBS at 37°C for 15 min to dissociate cells. Trypsin was neutralized with serum-containing medium. Cells were replated in medium without antibiotics in a 24-well plate at a density of 3 x 10^5^ cells/well and unprimed prior to infection as we have not observed differential inflammasome-dependent cytokine release in IECs during infection with priming (49). All Caco-2 knockout cell lines are described previously (49).

#### Cell culture of polarized Caco-2 cells

Polarized Caco-2 cells were grown on polycarbonate 3 μM pore size cell culture inserts (Corning 3415) in a 24 well plate. Inserts were coated with collagen coating solution containing 30 μg/ml collagen, 10 μg/ml fibronectin and 10 μg/ml BSA in DMEM and incubated for 3 hours. Caco-2 cells were then plated in growth medium containing Corning MITO+ serum extender (Fisher Scientific CB-50006) on inverted (for basolateral infection) or noninverted (for apical infection) inserts. After 24 hours, the growth medium was replaced with Corning enterocyte differentiation medium (Fisher Scientific 355357) with MITO+ serum extender. The media was replaced daily and following three days of incubation in differentiation media, the transepithelial electrical resistance was measured using an Epithelial Volt/Ohm (TEER) Meter (World Precision Instruments) to ensure that resistance was above 250 Ω.cm^2^ prior to infection. Infections were administered on the apical or basolateral side of cells as indicated.

#### Cell culture of THP-1 monocyte derived macrophages

THP-1 macrophages (TIB-202; American Type Culture Collection) were maintained in RPMI supplemented with 10% (vol/vol) heat-inactivated fetal bovine serum (FBS), 0.05 nM β-mercaptoethanol, 100 IU/mL penicillin and 100 μg/mL streptomycin. Two days prior to infection, THP-1 cells were replated in medium without antibiotics in a 48-well plate at a density of 2 x 10^5^ cells/well and incubated with phorbol 12-myristate 13-acetate (PMA) for 24 hours to allow differentiation into macrophages. Macrophages were primed with 100 ng/mL Pam3CSK4 (Invivogen) for 16 hours prior to bacterial infections in order to upregulate pro-IL-1β transcript levels.

#### Cell culture of primary human monocyte-derived macrophages (hMDMs)

Purified human monocytes from de-identified healthy human donors were obtained from the University of Pennsylvania Human Immunology Core. Monocytes were differentiated into macrophages by culturing in RPMI supplemented with 10% (vol/vol) heat-inactivated FBS, 2 mM L-glutamine, 100 IU/mL penicillin, 100 μg/ml streptomycin, and 50 ng/ml recombinant human M-CSF (Gemini Bio-Products) for 6 days. Two days prior to infection, adhered hMDMs were replated in media with 25 ng/ml human M-CSF lacking antibiotics at 1×10^5^ cells/well in a 48 well plate. hMDMs were then primed with 100 ng/ml Pam3CSK4 (Invivogen) for 16 hours prior to bacterial infection in order to upregulate pro-IL-1β transcript levels.

#### Cell culture of murine bone marrow derived macrophages (BMDMs)

Bone marrow cells were grown in RPMI containing L-cell supernatant, heat-inactivated FBS, penicillin and streptomycin for 8 days. One day prior to infection, differentiated BMDMs were replated into 24-well dishes in media lacking antibiotics at a density of 2 x 10^5^ cells/well.

### ELISAs

Supernatants harvested from infected cells were assayed using enzyme-linked immunosorbent assay (ELISA) kits for human IL-18 (R&D Systems) and IL-1β (BD Biosciences).

### LDH cytotoxicity assays

Supernatants harvested from infected cells were assayed for cytotoxicity by measuring loss of cellular membrane integrity via lactate dehydrogenase (LDH) assay. LDH release was quantified using an LDH Cytotoxicity Detection Kit (Clontech) according to the manufacturer’s instructions and normalized to mock-infected (min cytotoxicity) and 2% triton-treated cells (max cytotoxicity)

### Immunoblot analysis

Cells were replated and infected on serum-free medium to collect supernatant samples. Supernatant samples were centrifuged at 200 x *g* to pellet any cell debris and treated with trichloroacetic acid (TCA) (25 μL TCA per 500 μL supernatant) overnight at 4°C. The following day, TCA-treated samples were centrifuged at max speed (15,871 x *g*) for 15 min at 4°C and washed with ice-cold acetone. TCA-precipitated supernatant samples and cell lysates were resuspended in 1 x SDS-PAGE sample buffer and boiled for 5 min. Samples were separated by SDS-PAGE on a 12% (vol/vol) acrylamide gel and transferred to polyvinylidene difluoride (PVDF) Immobilon-P membranes (Millipore). Primary antibodies specific for human IL-18 (MLB International PM014), IL-1β (R&D Systems MAB201), β-actin (4967L; Cell Signaling) and GSDMD (G7422 Sigma-Aldrich) and horseradish peroxidase (HRP)-conjugated secondary antibodies anti-rabbit IgG (7074S; Cell Signaling) and anti-mouse IgG (7076S; Cell Signaling) were used. Enhanced chemiluminescence (ECL) Western blotting substrate or SuperSignal West Femto (Pierce Thermo Scientific) HRP substrate were used for detection.

### Inhibitor and antibody blocking experiments

Cells were treated 1 h prior to infection at the indicated concentrations of the following inhibitors: 10 μM MCC950 (Sigma-Aldrich; PZ0280), 20 μM pan-caspase inhibitor Z-VAD(Ome)-FMK (SM Biochemicals; SMFMK001), 20 μM caspase-1 inhibitor Ac-YVAD-cmk (Sigma-Aldrich; SML0429), 30 μM disulfiram (Sigma), 10 μM cytochalasin D (Sigma), and 25 μM Akt inhibitor VIII (EMD Millipore). Cells were treated 1 h prior to infection with a 1:50 dilution of monoclonal antibody 1987 clone P4C10 (EMD Millipore) directed against β1 integrins.

### siRNA-mediated gene knockdown

*CASP5* (S2417), *CASP8* (S2427), *ITGA5* (S7549) and two Silencer Select negative-control siRNAs (Silencer Select negative control no. 1 and no. 2 siRNA) were purchased from Ambion (Life Technologies). Three days before infection, 30 nM siRNA was transfected into Caco-2 cells using Lipofectamine RNAiMAX transfection reagent (Thermo Fisher Scientific) following the manufacturer’s protocol.

### Quantitative RT-PCR analysis

RNA was isolated using the RNeasy Plus Mini Kit (Qiagen) following the manufacturer’s instructions. Cells were lysed in 350 μL RLT buffer with β-mercaptoethanol and centrifuged through a QIAshredder spin column (Qiagen). cDNA was synthesized from isolated RNA using SuperScript II Reverse Transcriptase (Invitrogen) following the manufacturer’s instructions. Quantitative PCR was conducted with the CFX96 real-time system from Bio-Rad using the SsoFast EvaGreen Supermix with Low ROX (Bio-Rad). For analysis, mRNA levels of siRNA-treated cells were normalized to housekeeping gene *HPRT* and control siRNA-treated cells using the 2^-ϕλϕλCT^ (cycle threshold) method to calculate knockdown efficiency (129). The following primers were used:

*CASP5* forward: TTCAACACCACATAACGTGTCC

*CASP5* reverse: GTCAAGGTTGCTCGTTCTATGG

*CASP8* forward: GTTGTGTGGGGTAATGACAATCT

*Casp8* reverse: TCAAAGGTCGTGGTCAAAGCC

*ITGA5* forward: GGCTTCAACTTAGACGCGGAG

*ITGA5* reverse: TGGCTGGTATTAGCCTTGGGT

*HPRT* forward: CCTGGCGTCGTGATTAGTGAT

*HPRT* reverse: AGACGTTCAGTCCTGTCCATAA

### Bacterial uptake enumeration with colony forming units (CFUs)

Cells were infected with indicated strains of *Yersinia* at an MOI of 20. 1 hpi, cells were treated with 100 μg/mL of gentamicin to kill extracellular bacteria. 2 hpi the supernatants were aspirated and cells were lysed with PBS containing 0.5% Triton to collect intracellular bacteria. Harvested bacteria were serially diluted in PBS and plated on LB agar plates containing 2 μg/mL Irgasan. Plates were incubated at 28°C for two days and CFUs were counted.

### Fluorescence microscopy of intracellular *Yersinia*

One day before infection 2 x 10^5^ cells/well were plated on glass coverslips in a 24-well plate. Cells were infected with indicated strains of *Yersinia* constitutively expressing GFP at an MOI of 20. At 2hpi, cells were washed 2 times with PBS, fixed with 4% paraformaldehyde for 10 min at 37°C and stored overnight at 4°C in PBS. The following day, cells were blocked for 30 min at room temperature in blocking solution containing 1% BSA in PBS and incubated for 1 h at room temperature in blocking solution with the polyclonal anti-*Yersinia* antibody SB349 diluted 1:1000 (kindly provided by Dr. James Bliska) (102). AF594-conjugated goat anti-Rabbit IgG antibody (A-11012 Thermo Fisher Scientific) was diluted 1:500 in blocking solution was added to cells and incubated for 45 min at room temperature. Cells were mounted on glass slides with DAPI mounting medium (Sigma Fluoroshield). Coverslips were imaged on an inverted fluorescence microscope (IX81; Olympus) and images were collected using a high-resolution charge-coupled-devise camera (FAST1394; QImaging) at a magnification of 60x. Images were analyzed and presented using SlideBook (version 5.0) software (Intelligent Imaging Innovations, Inc.). %intracellular bacteria were scored in unblinded fashion by counting 20 captures per coverslip for coverslips across independent triplicate experiments.

### Statistical analysis

Prism 9.4.1 (GraphPad Software) was utilized for the graphing of data and all statistical analyses. Statistical significance for experiments were determined using the appropriate test and are indicated in each figure legend. Differences were considered statistically significant if the *p* value was <0.05.

## Data availability

All data are included in the manuscript and supplemental material. Bacterial strains available upon request

## Acknowledgments

We thank members of the Shin and Brodsky laboratories for helpful scientific discussions. We thank Dr. James Bliska, Dr. Joan Mecsas, and Dr. Petra Dersch for generously providing plasmids and Yop mutant strains.

We thank the Human Immunology Core of the Penn Center for AIDS Research and Abramson Cancer Center for providing purified primary human monocytes.

This work is supported by NIH/NIAID grants AI151476, AI118861, AI123243 (S.S.), AI128630, AI163596, and AI139102 (I.E.B). S.S. and I.E.B. are both recipients of the Burroughs-Welcome Fund Investigators in the Pathogenesis of Infectious Disease Award. J.Z. is a recipient of the NIH/NIAID Microbial Pathogenesis and Genomics training grant 5T32AI141393-03.

