## Supplemental figures for "*Yersinia* Type III-Secreted Effectors Evade the Caspase-4 Inflammasome in Human Cells"

Table S1. *Yersinia* strains used in this study

| Strain name | Relevant characteristics | Reference or Source |
|---|---|---|
| IP2666 (WT *Yptb*) | Wild-type, $pYV^{+}$, naturally *yopT*$^{-}$ | (128) |
| *Δ6 Yptb* | *yopEHJMKO*$^{-}$ | (79) |
| Δ*yopE Yptb* | *yopE-* | (102) |
| Δ*yopH Yptb* | *yopH-* | (105) |
| Δ*yopM Yptb* | *yopM-* | This study and (40) |
| Δ*yopJ Yptb* | *yopJ-* | (79) |
| Δ*yopK Yptb* | *yopK-* | (45) |
| Δ*yopO Yptb* | *yopO-* | (130) |
| Δ*yopEH Yptb* | *yopEH-* | (105) |
| Δ*yopEK Yptb* | *yopEK-* | This study and (131) |
| Δ*yopHK Yptb* | *yopHK-* | This study and (131) |
| Δ*yopEHK Yptb* | *yopEHK-* | This study and (131) |
| Δ*yopEH*Δ*inv Yptb* | *yopEH-* invasin- | This study and (119) |
| Δ*yopEH*Δ*yadA Yptb* | *yopEH-* YadA- | This study and (70) |
| Δ*yopEH*Δ*inv*Δ*yadA Yptb* | *yopEH-* invasin- YadA- | This study and (70, 119) |
| *Δ6Δinv Yptb* | *yopEHJMKO-* invasin- | This study and (119) |
| *Δ6ΔyadA Yptb* | *yopEHJMKO-* YadA- | This study and (70) |
| *Δ6ΔinvΔyadA Yptb* | *yopEHJMKO-* invasin- YadA- | This study and (70, 119) |

Figure S1 (related to Fig. 1). *Yersinia* Yops evade inflammasome activation. (A, E) THP-1 macrophages were infected with PBS (Mock) or WT *Yptb* or Δ*6 Yptb*. (A) Percent cytotoxicity normalized to cells treated with 2% triton and (E) release of IL-1β into the supernatant was measured by ELISA at 6hpi. (B) Polarized Caco-2 cells were infected with PBS or WT *Yptb* or Δ*6 Yptb*. Release of IL-18 into the apical and basolateral supernatant was measured by ELISA at 6hpi. (C) WT Caco-2 cells were infected with the indicated MOI and strain of *Yptb*. IL-18 release was measured by ELISA at 6hpi. (D) WT Caco-2 cells were infected with PBS (Mock) or WT *Yptb* or Δ*6 Yptb*. Release of IL-1β into the supernatant was measured by ELISA at 6hpi. ** $p < 0.01$, *** $p < 0.001$, **** $p < 0.0001$ by one-way ANOVA. Shown are averages and error bars representing the standard deviation from at least three pooled experiments.

Figure S2 (related to Fig. 2). Caspase-1 and caspase-8 are not absolutely required for Δ*6 Yptb*-induced inflammasome activation in human IECs. (A) WT Caco-2 cells or two independent single cell clones of *CASP1*$^{-/-}$ Caco-2 cells were infected with PBS (Mock), WT *Yptb*, or Δ*6 Yptb*. (B) One hour prior to infection, WT Caco-2 cells were treated with 20 μM YVAD or DMSO as a vehicle control. Cells were then infected with PBS (Mock), WT *Yptb* or Δ*6 Yptb*. (A, B) IL-18 release into the supernatant was measured by ELISA at 6hpi. (C) Knockdown efficiency of siRNA targeting *CASP8* in WT Caco-2 cells was measured by qRT-PCR and normalized to housekeeping gene *HPRT* and calculated relative to control-siRNA-treated cells. (D) WT Caco-2 cells were treated with siRNA targeting a control scrambled siRNA or siRNA targeting *CASP8* for 72 hours. (E) One hour prior to infection WT Caco-2 cells were treated with 20 μM IETD or DMSO as a vehicle control. (D, E) Cells were infected with PBS (Mock), WT *Yptb* or Δ*6 Yptb*. IL-18 release into the supernatant was measured at 6hpi. (F, G) WT or two independent single cell clones of *CASP1*$^{-/-}$ Caco-2 cells were treated with siRNA targeting a control scrambled siRNA or siRNA targeting *CASP8* for 72 hours. (F) IL-18 release into the supernatant was measured at

6hpi. (G) Knockdown efficiency of siRNA targeting *CASP8* in WT Caco-2 cells was measured by qRT-PCR and normalized to housekeeping gene *HPRT* and calculated relative to control-siRNA-treated cells. *** $p < 0.001$ by two-way ANOVA. (A-E) Shown are averages and error bars representing the standard deviation from at least three pooled experiments. (F, G) Error bars represent standard deviation of triplicate wells and are representative of two independent experiments.

Figure S3 (related to Fig. 2). Caspase-4 contributes to $\Delta 6$ *Yptb*-induced inflammasome activation in human macrophages. (A) WT or two independent single cell clones of *CASP4*$^{-/-}$ THP-1 macrophages were infected with PBS (Mock), WT *Yptb* or $\Delta 6$ *Yptb*. (B) One hour prior to infection WT THP-1 macrophages were treated with 20 $\mu$M ZVAD or DMSO as a vehicle control. (A, B) IL-1$\beta$ release into the supernatant was measured at 6hpi. (C) Knockdown efficiency of siRNA targeting *CASP5* in WT Caco-2 cells was measured by qRT-PCR, normalized to *HPRT* and calculated relative to control-siRNA-treated cells. (D) WT Caco-2 cells were treated with siRNA targeting a control scrambled siRNA or siRNA targeting *CASP5* for 72 hours. IL-18 release was measured at 6hpi. ** $p < 0.01$, *** $p < 0.001$ by two-way ANOVA. Shown are averages and error bars representing the standard deviation from at least three pooled experiments.

Figure S4 (related to Fig. 3). NLRP3, NAIP/NLRC4 and ASC are dispensable for $\Delta 6$ *Yptb*-induced inflammasome activation in human IECs. (A) One hour prior to infection cells were treated with 10 $\mu$M MCC950 or DMSO as a vehicle control. Cells were then infected with PBS (Mock) WT *Yptb* or $\Delta 6$ *Yptb.* (B, C) WT or two independent single cell clones of (B) *NAIP*$^{-/-}$ or (C) *PYCARD*$^{-/-}$ Caco-2 cells were infected with PBS (Mock), WT *Yptb*, or $\Delta 6$ *Yptb*. (A, B, C) IL-18 release was measured in supernatants at 6hpi. Shown are averages and error bars representing the standard deviation from at least three pooled experiments.

Figure S5 (related to Fig. 5). YopE and YopH block internalization in human cells. (A) WT Caco-2 cells or (B) WT THP-1 macrophages were infected with the indicated strain of *Yptb* at an MOI of 20 and lysed at 2 hpi. Bacteria were plated on *Yersinia*-selective agar to enumerate CFUs. (C, D) WT Caco-2 cells were seeded on glass coverslips and infected with indicated strain of *Yptb* expressing GFP at an MOI of 20. Cells were fixed at 2hpi and stained for extracellular *Yptb* (Red) and DAPI to label DNA (blue). (C) Representative images are shown. Scale bar represents 10 $\mu$m. (D) Proportion of total bacteria that was intracellular (green only) was scored unblinded by fluorescence microscopy. Shown are averages and error bars of coverslips from independent experiments with 20 fields scored per coverslip. (E, F) One hour prior to infection, WT Caco-2 cells were treated with 10 $\mu$M cytochalasin D or DMSO as a vehicle control. Cells were then infected with PBS (Mock) or indicated strain of *Yptb* at an MOI of 20 (E) or 60 (F). (E) Cells were lysed at 2hpi and bacteria were plated on *Yersinia*-selective agar to calculate CFUs. (F) Release of IL-18 into the supernatant was measured at 6hpi. * $p < 0.05$, *** $p < 0.001$, **** $p < 0.0001$ by one-way ANOVA (A, B, D) or two-way ANOVA (E, F). Shown are averages and error bars representing the standard deviation from at least three pooled experiments.

Figure S6 (related to Fig. 5). (A, B) One hour prior to infection, WT Caco-2 cells were treated with 25 $\mu$M Akt inhibitor VIII or DMSO as a vehicle control. Cells were then infected with PBS (Mock) or indicated strain of *Yptb* at an MOI of 20 (A) or 60 (B). (A) Cells were lysed at 2hpi and bacteria were plated on *Yersinia*-selected agar to calculate CFUs. (B) Release of IL-18 into the supernatant was measured at 6hpi. (C, E) WT Caco-2 cells were treated with a control scrambled siRNA or siRNA targeting *ITGA5* for 72 hours. Cells were then infected with PBS (Mock) or indicated strain of *Yptb* at an MOI of 20 (C) or 60 (D). (C) Cells were lysed at 2hpi and bacteria were plated on *Yersinia*-selected agar to calculate CFUs. (D) Release of IL-18 into the supernatant was measured at 6hpi. (E) Knockdown efficiency of siRNA targeting *ITGA5* in WT Caco-2 cells was measured by qRT-PCR and normalized to housekeeping gene *HPRT* and calculated relative to control-siRNA-treated cells. (F, G) WT Caco-2 cells were infected with PBS (Mock) or indicated strain of *Yptb* at an MOI of 20 (F) or 60 (G). (F) Cells were lysed at 2hpi and bacteria were plated on *Yersinia*-selected agar to calculate CFUs. (G) Release of IL-18 into the supernatant was measured at 6hpi. Shown are averages and error bars representing the standard deviation from at least three pooled experiments.

Figure S1

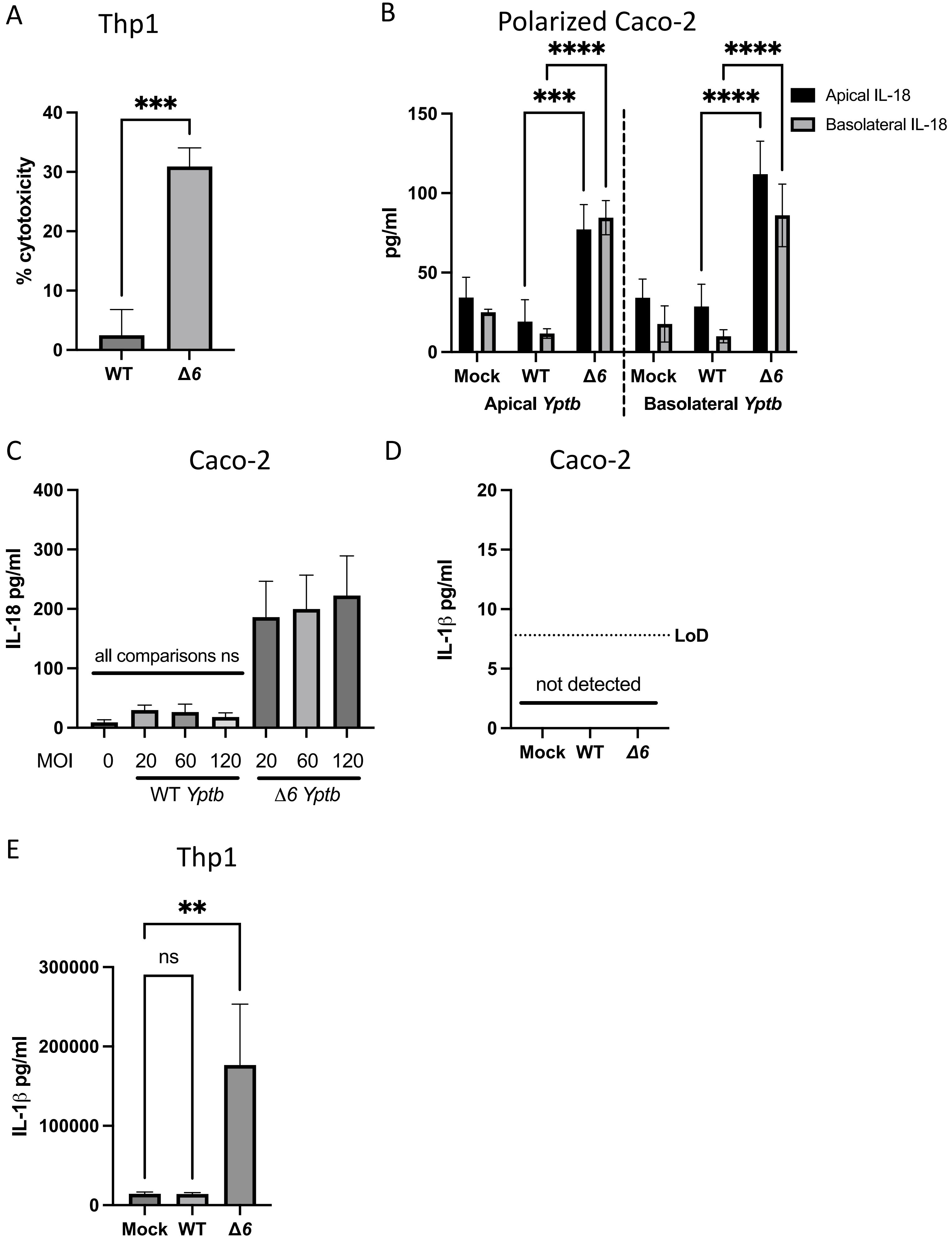

A
Thp1
***
40
30
20
10
0
% cytotoxicity
WT
Δ6
B
Polarized Caco-2
****
***
****
****
Apical IL-18
Basolateral IL-18
150
100
50
0
pg/ml
Mock
WT
Δ6
Mock
WT
Δ6
Apical Yptb
Basolateral Yptb
C
Caco-2
400
300
200
100
0
IL-18 pg/ml
all comparisons ns
MOI
0
20
60
120
20
60
120
WT Yptb
Δ6 Yptb
D
Caco-2
20
15
10
5
0
IL-1β pg/ml
LoD
not detected
Mock
WT
Δ6
E
Thp1
**
ns
300000
200000
100000
0
IL-1β pg/ml
Mock
WT
Δ6

**Figure S2**

A

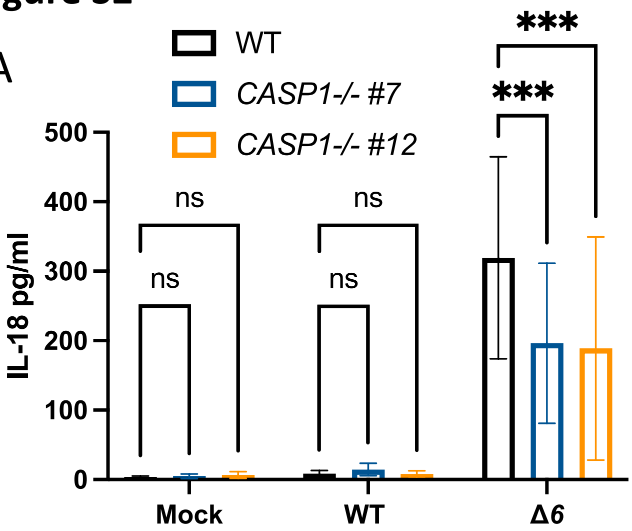


B

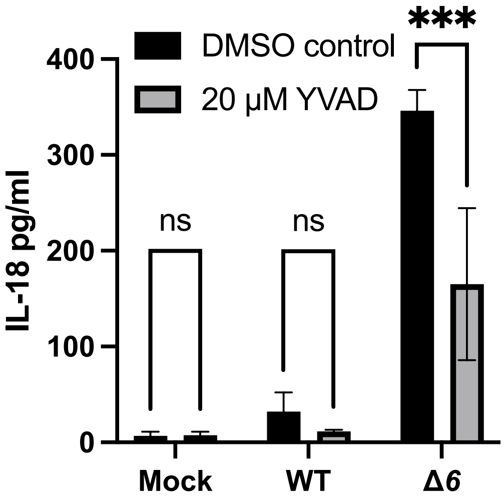


C

Average knockdown efficiency

| | *CASP8* |
|---|---|
| WT | 66% |

D

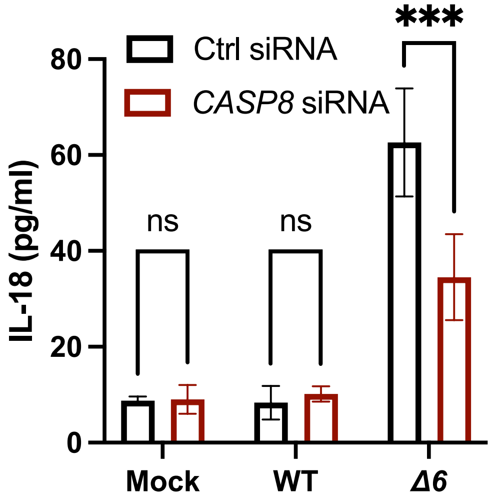


E

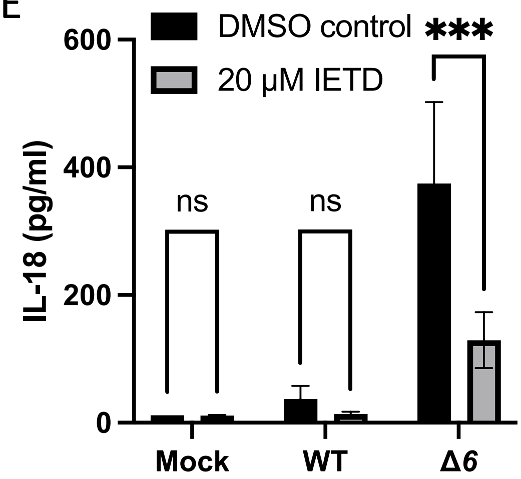


F

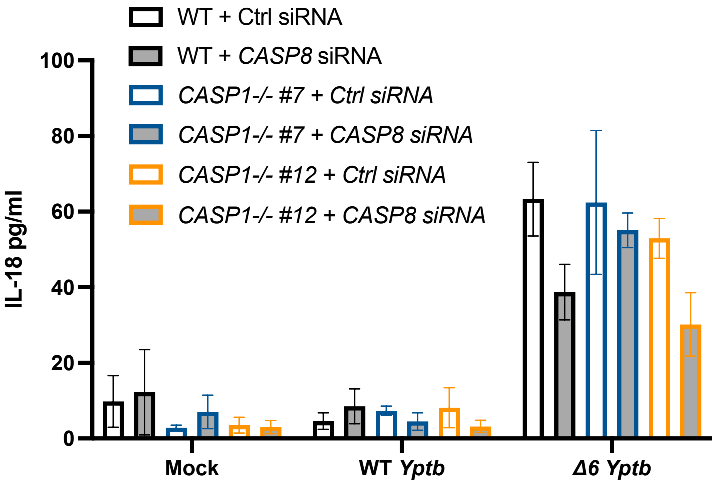


G

Average knockdown efficiency

| | *CASP8* |
|---|---|
| WT | 57% |
| *CASP1-/- #7* | 68% |
| *CASP1-/- #12* | 74% |

**Figure S3**

A

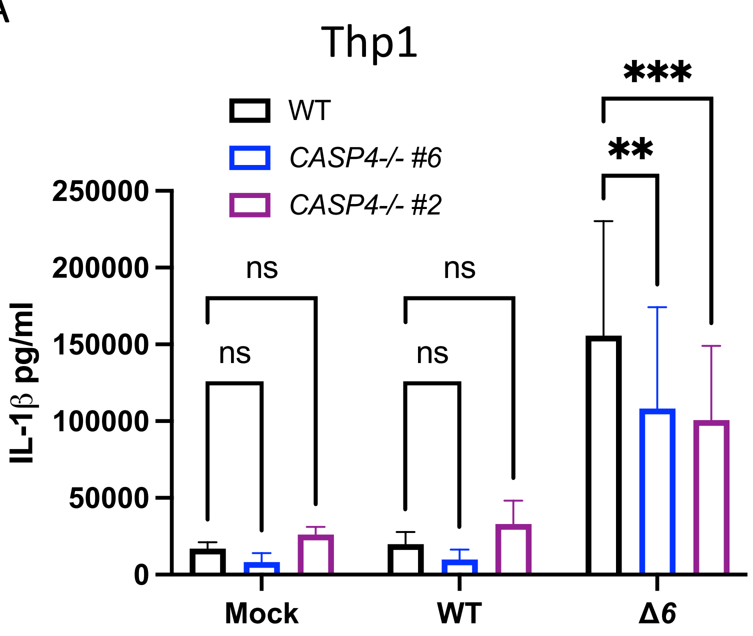


B

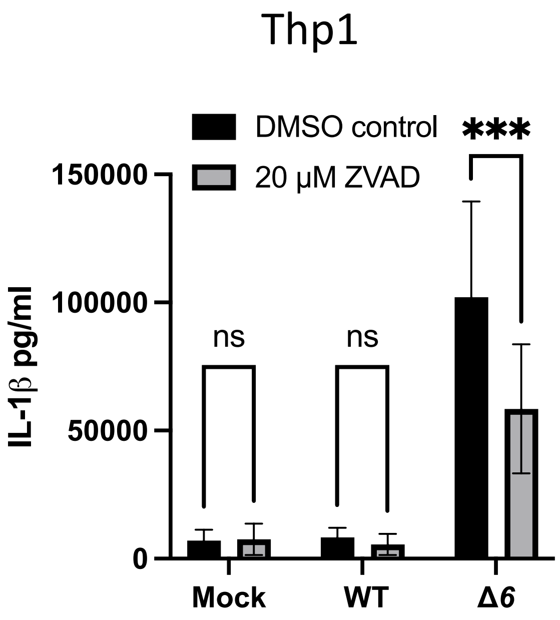


C

Caco-2

Average knockdown efficiency

| | *CASP5* |
|---|---|
| WT Caco-2s | 75% |

D

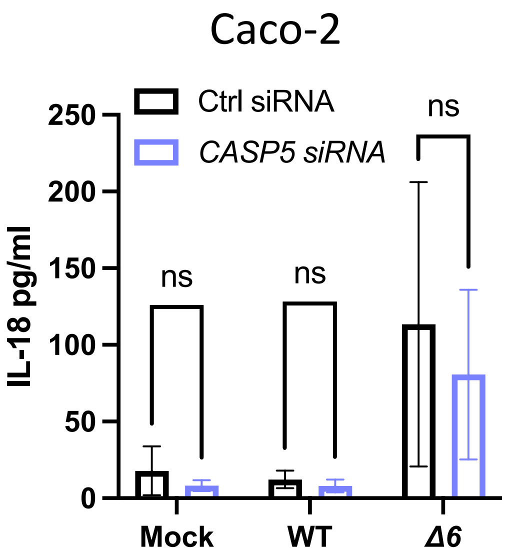

**Figure S4**

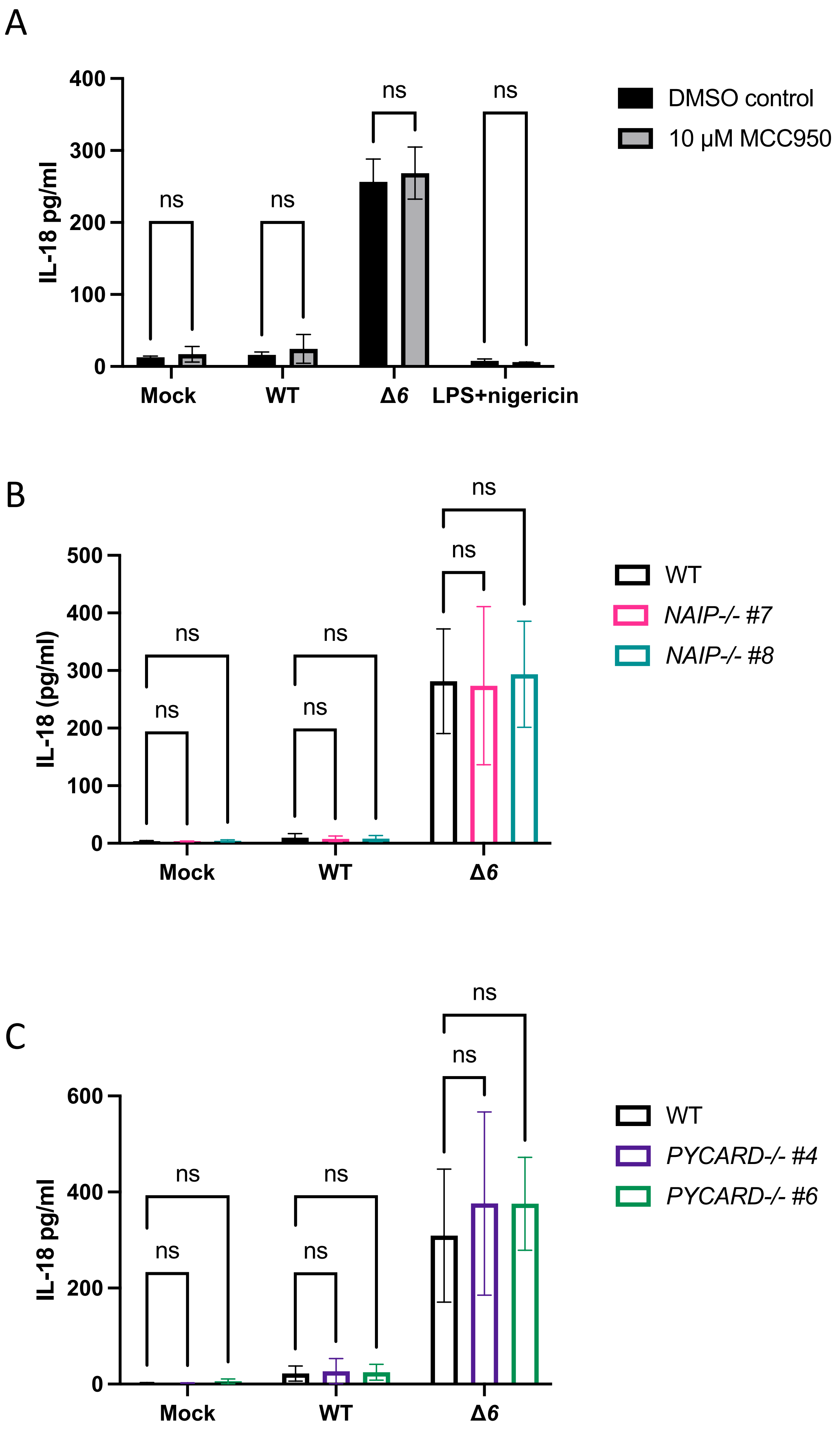
A
IL-18 pg/ml
400
300
200
100
0
ns
Mock
WT
Δ6
LPS+nigericin
DMSO control
10 µM MCC950
B
IL-18 (pg/ml)
500
400
300
200
100
0
ns
Mock
WT
Δ6
WT
NAIP-/- #7
NAIP-/- #8
C
IL-18 pg/ml
600
400
200
0
ns
Mock
WT
Δ6
WT
PYCARD-/- #4
PYCARD-/- #6

**Figure S5**

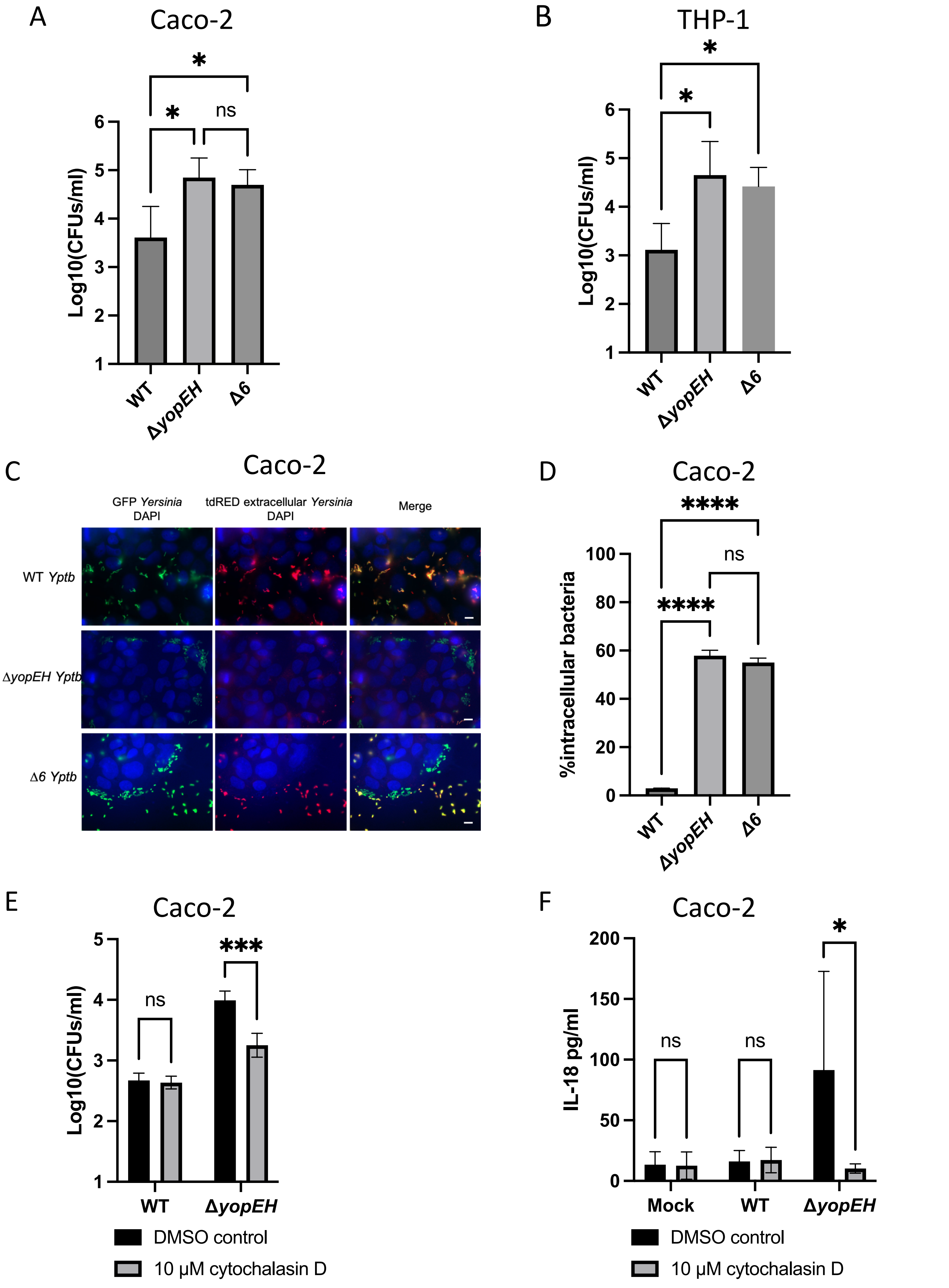

A
Caco-2
*
*
ns
Log10(CFUs/ml)
6
5
4
3
2
1
WT
ΔyopEH
Δ6
B
THP-1
*
*
Log10(CFUs/ml)
6
5
4
3
2
1
WT
ΔyopEH
Δ6
C
Caco-2
GFP Yersinia
DAPI
tdRED extracellular Yersinia
DAPI
Merge
WT Yptb
ΔyopEH Yptb
Δ6 Yptb
D
Caco-2
****
ns
****
%intracellular bacteria
100
80
60
40
20
0
WT
ΔyopEH
Δ6
E
Caco-2
ns
***
Log10(CFUs/ml)
5
4
3
2
1
WT
ΔyopEH
DMSO control
10 μM cytochalasin D
F
Caco-2
ns
ns
*
IL-18 pg/ml
200
150
100
50
0
Mock
WT
ΔyopEH
DMSO control
10 μM cytochalasin D

**Figure S6**

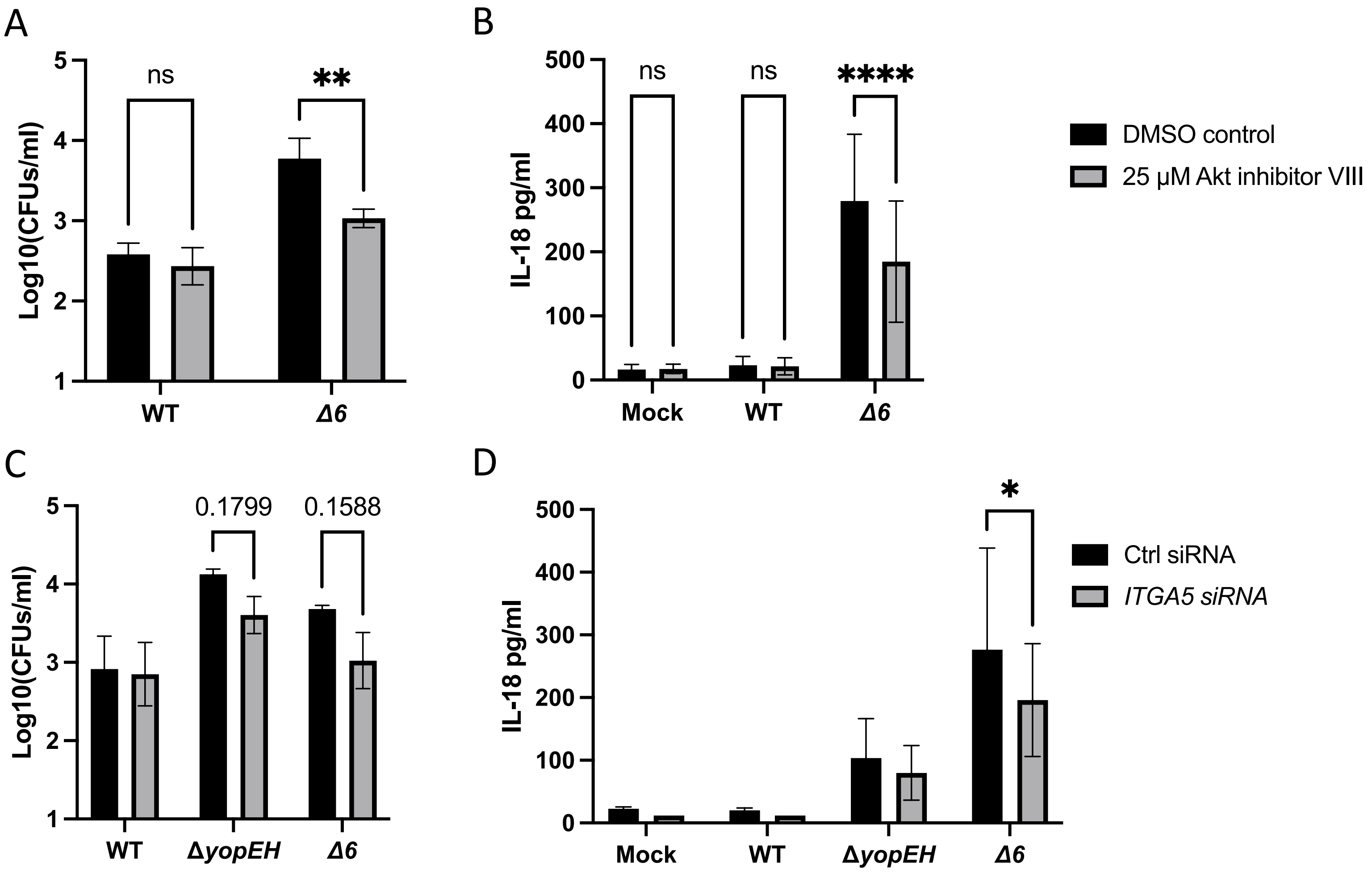


E

Average knockdown efficiency

| | *ITGA5* |
|---|---|
| WT | 65% |

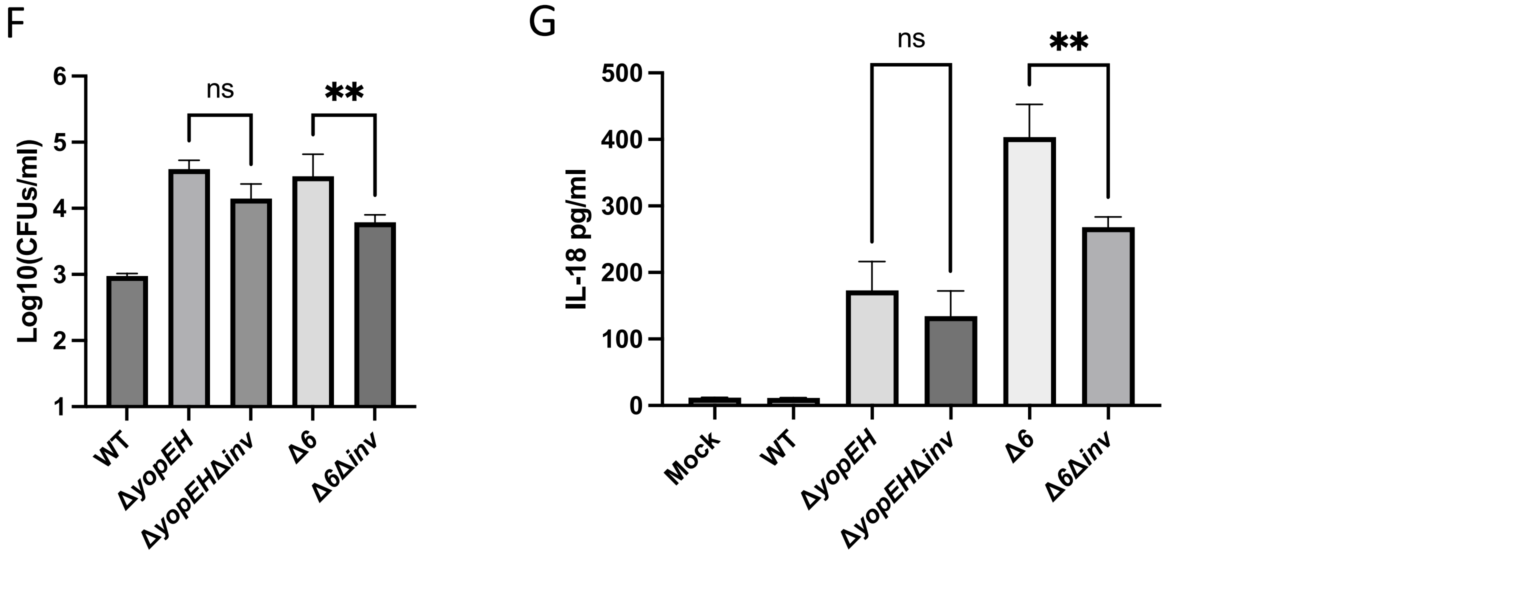
